## Supplemental Text for "Integrating healthcare and research genetic data empowers the discovery of 49 novel developmental disorders"

#### Supplemental Contents

**DDD Study consortium authors** 3

**GeneDx Data Sharing** 5

**Sample collection and individual QC** 6
- DDD 6
- GeneDx 6
- Radboud University Medical Center 7

**Definition of diagnostic lists** 8

**Joint quality control of dataset** 10
- De novo mutation filtering 10
- Duplicate samples 13
- Removing variants from siblings 13

**Comparing datasets** 14

**Confirming variants in novel genes** 15

**Enrichment framework** 17
- DeNovoWEST 17
  - Determination of Weights 18
  - De Novo Clustering Test 20
  - Combining Tests 20
  - Independent Hypothesis Weighting Multiple Testing Correction 20
- Comparing DeNovoWEST to previous enrichment test 22
- Synonymous-only enrichment test 23
- Analysis of undiagnosed cases 24
- Withholding the 10 most mutated genes from weight creation 25

**Evaluating potential cryptic splice site variants** 26

**Estimating the fraction of cases explained by de novo mutations** 28
- Splitting by cohort 28
- Splitting by sex 28

**Comparing the new and known genes** 30
- Phenotypic similarity between new and known genes 30
- Functional similarity between new and known genes 30
- Evaluating potentially smaller mutational targets in the novel genes 33
- Comparison of constraint metrics 33
- Clustering of PTVs at the end of transcripts 34

**Protein domain analysis** 35
- Annotation of transcripts, protein positions, and protein domains to genomic coordinates 35
- Counting variants by type and correcting for region-specific bias 35

Computing variant enrichment in protein domains 37

**Recurrent Mutations** 40

**Overlap with somatic driver genes and mutations** 41

**DNM burden in non-significant genes** 43

**Impact of pre/perinatal death on power** 44

**Penetrance** 46

**Power analysis** 47

**Downsampling** 48

**Modelling remaining DNM burden** 49

Model for PTV DNM burden 50

Model for missense DNM burden 51

**Expression in fetal brain** 52

### DDD Study consortium authors

*Aberdeen*
Silvia Borras, Caroline Clark, John Dean, Zosia Miedzybrodzka, Alison Ross, Stephen Tennant

*Belfast*
Tabib Dabir, Deirdre Donnelly, Mervyn Humphreys, Alex Magee, Vivienne McConnell, Shane McKee, Susan McNerlan, Patrick J. Morrison, Gillian Rea, Fiona Stewart

*Birmingham*
Trevor Cole, Nicola Cooper, Lisa Cooper-Charles, Helen Cox, Lily Islam, Joanna Jarvis, Rebecca Keelagher, Derek Lim, Dominic McMullan, Jenny Morton, Swati Naik, Mary O'Driscoll, Kai-Ren Ong, Deborah Osio, Nicola Ragge, Sarah Turton, Julie Vogt, Denise Williams

*Bristol*
Simon Bodek, Alan Donaldson, Alison Hills, Karen Low, Ruth Newbury-Ecob, Andrew M. Norman, Eileen Roberts, Ingrid Scurr, Sarah Smithson, Madeleine Tooley

*Cambridge*
Steve Abbs, Ruth Armstrong, Carolyn Dunn, Simon Holden, Soo-Mi Park, Joan Paterson, Lucy Raymond, Evan Reid, Richard Sandford, Ingrid Simonic, Marc Tischkowitz, Geoff Woods

*Dublin*
Lisa Bradley, Joanne Comerford, Andrew Green, Sally Lynch, Shirley McQuaid, Brendan Mullaney

*Dundee*
Jonathan Berg, David Goudie, Eleni Mavrak, Joanne McLean, Catherine McWilliam, Eleanor Reavey

*Edinburgh*
Tara Azam, Elaine Cleary, Andrew Jackson, Wayne Lam, Anne Lampe, David Moore, Mary Porteous

*Exeter*
Emma Baple, Júlia Baptista, Carole Brewer, Bruce Castle, Emma Kivuva, Martina Owens, Julia Rankin, Charles Shaw-Smith, Claire Turner, Peter Turnpenny, Carolyn Tysoe

*Glasgow*
Therese Bradley, Rosemarie Davidson, Carol Gardiner, Shelagh Joss, Esther Kinning, Cheryl Longman, Ruth McGowan, Victoria Murday, Daniela Pilz, Edward Tobias, Margo Whiteford, Nicola Williams

*GOSH, London*
Angela Barnicoat, Emma Clement, Francesca Faravelli, Jane Hurst, Lucy Jenkins, Wendy Jones, V.K.Ajith Kumar, Melissa Lees, Sam Loughlin, Alison Male, Deborah Morrogh, Elisabeth Rosser, Richard Scott, Louise Wilson

*Guy's, London*

Ana Beleza, Charu Deshpande, Frances Flinter, Muriel Holder, Melita Irving, Louise Izatt, Dragana Josifova, Shehla Mohammed, Aneta Molenda, Leema Robert, Wendy Roworth, Deborah Ruddy, Mina Ryten, Shu Yau

*Leeds*
Christopher Bennett, Moira Blyth, Jennifer Campbell, Andrea Coates, Angus Dobbie, Sarah Hewitt, Emma Hobson, Eilidh Jackson, Rosalyn Jewell, Alison Kraus, Katrina Prescott, Eamonn Sheridan, Jenny Thomson

*Leicester and Nottingham*
Kirsty Bradshaw, Abhijit Dixit, Jacqueline Eason, Rebecca Haines, Rachel Harrison, Stacey Mutch, Ajoy Sarkar, Claire Searle, Nora Shannon, Abid Sharif, Mohnish Suri, Pradeep Vasudevan

*Liverpool*
Natalie Canham, Ian Ellis, Lynn Greenhalgh, Emma Howard, Victoria Stinton, Andrew Swale, Astrid Weber

*Manchester*
Siddharth Banka, Catherine Breen, Tracy Briggs, Emma Burkitt-Wright, Kate Chandler, Jill Clayton-Smith, Dian Donnai, Sofia Douzgou, Lorraine Gaunt, Elizabeth Jones, Bronwyn Kerr, Claire Langley, Kay Metcalfe, Audrey Smith, Ronnie Wright

*Newcastle*
David Bourn, John Burn, Richard Fisher, Steve Hellens, Alex Henderson, Tara Montgomery, Miranda Splitt, Volker Straub, Michael Wright, Simon Zwolinski

*NW Thames, London*
Zoe Allen, Birgitta Bernhard, Angela Brady, Claire Brooks, Louise Busby, Virginia Clowes, Neeti Ghali, Susan Holder, Rita Ibitoye, Emma Wakeling

*Oxford*
Edward Blair, Jenny Carmichael, Deirdre Cilliers, Susan Clasper, Richard Gibbons, Usha Kini, Tracy Lester, Andrea Nemeth, Joanna Poulton, Sue Price, Debbie Shears, Helen Stewart, Andrew Wilkie

*Sheffield*
Shadi Albaba, Duncan Baker, Meena Balasubramanian, Diana Johnson, Michael Parker, Oliver Quarrell, Alison Stewart, Josh Willoughby

*St. George’s, London*
Charlene Crosby, Frances Elmslie, Tessa Homfray, Huilin Jin, Nayana Lahiri, Sahar Mansour, Karen Marks, Meriel McEntagart, Anand Saggar, Kate Tatton-Brown

*Wales*
Rachel Butler, Angus Clarke, Sian Corrin, Andrew Fry, Arveen Kamath, Emma McCann, Hood Mugalaasi, Caroline Pottinger, Annie Procter, Julian Sampson, Francis Sansbury, Vinod Varghese

*Wessex*
Diana Baralle, Alison Callaway, Emma J. Cassidy, Stacey Daniels, Andrew Douglas, Nicola Foulds, David Hunt, Mira Kharbanda, Katherine Lachlan, Catherine Mercer, Lucy Side, I. Karen Temple, Diana Wellesley

### GeneDx Data Sharing

The American College of Genetics and Genomics (ACMG) position statement on genomic data sharing helps inform GeneDx's philosophy regarding data use: "To ensure that our patients receive the most informed care possible, the American College of Medical Genetics and Genomics advocates for extensive sharing of laboratory and clinical data from individuals who have undergone genomic testing. Information that underpins health-care service delivery should be treated neither as intellectual property nor as a trade secret when other patients may benefit from the knowledge being widely available" [1]. With this in mind, as part of its clinical testing program, per patient consent and without sharing any non-deidentified data, and as described on the website (https://www.genedx.com/all-forms/use-of-specimen-and-genetic-information/). GeneDx uses resources such as GeneMatcher[1,2] (https://genematcher.org) in order to help identify evidence that variants in a given gene may contribute to an observed phenotype. Similarly, the endeavor described in this work allows GeneDx analysts and scientists to identify affected patients in other cohorts, which translates to the ability to better clinically interpret variants in specific genes of interest and provide reports to referring clinicians. This process is also useful for updating variants from those binned as candidate genes to more clearly causative disease explanations; GeneDx has updated over 200 clinical reports regarding such genes since 2000, affecting more than 3,000 individual cases.

### Sample collection and individual QC

#### DDD

Patients with severe, undiagnosed developmental disorders were recruited from 24 regional genetics services within the United Kingdom National Health Service and the Republic of Ireland. Families gave informed consent to participate, and the study was approved by the UK Research Ethics Committee (10/H0305/83 granted by the Cambridge South Research Ethics Committee, and GEN/284/12 granted by the Republic of Ireland Research Ethics Committee). Additional details on sample collection, exome sequencing, alignment, variant calling (inherited and *de novo*) and variant annotation have been described previously[3]. These analyses involve 9,858 trios from 9,307 families, a subset of whom have been analyzed in previous publications[3,4].

#### GeneDx

Patients were referred to GeneDx for clinical whole-exome sequencing for diagnosis of suspected Mendelian disorders as previously described[5]. Patient medical records were abstracted into HPO terms using Neji concept recognition[6] with manual review by laboratory genetic counselors or clinicians. Patients were selected for inclusion in this study based on having one or more HPO phenotypes overlapping the inclusion criteria for the DDD study[4]. The study was conducted in accordance with all guidelines set forth by the Western Institutional Review Board, Puyallup, WA (WIRB 20162523). Informed consent for genetic testing was obtained from all individuals undergoing testing, and WIRB waived authorization for use of de-identified aggregate data. Individuals or institutions who opted out of this type of data use were excluded.

Data was sequenced and aligned as previously described[5] with either SureSelect Human All Exon v4 (Agilent Technologies, Santa Clara, CA), Clinical Research Exome (Agilent Technologies, Santa Clara, CA), or xGen Exome Research Panel v1.0 (IDT, Coralville, IA) and sequenced with either 2x100 or 2x150bp reads on HiSeq 2000, 2500, 4000, or NovaSeq 6000 (Illumina, San Diego, CA). Alignment BAM files were then converted to CRAM format with Samtools version 1.3.1 and indexed. Individual GVCF files were called with GATK v3.7-0 HaplotypeCaller[7,8] in GVCF mode by restricting output regions to plus/minus 50bp of the RefGene primary coding regions. Single-sample GVCF files were then combined into multi-sample GVCF files with each combined file contained 200 samples. These multi-sample GVCF files were then joint-genotyped using GATK GenotypeGVCFs. The cohort of 18,789 trios was joint-genotyped in two separate batches, one with 10,138 trios and the other 8651 trios. GATK VariantRecalibrator (VQSR) was applied for both SNPs and INDELs, with known SNPs from 1000 Genomes phase 1 high confidence set and “gold standard” INDELs from Mills et al[9].

Variants in VQSR VCF files were annotated with Ensembl Variant Effect Predictor (VEP)[10] using RefSeq transcripts. The transcript with the most severe consequence was selected, and all associated VEP annotations were based on the predicted effect of the variant on that particular transcript. Variants called in the proband and not in the parents were selected as potential *de novo* mutations. Filtering of these *de novo* mutations is described below.

#### Radboud University Medical Center

The Department of Human Genetics from the Radboud University Medical Center (RUMC) is a tertiary referral centre for clinical genetics. Approximately 350 individuals with unexplained intellectual disability (ID) are referred annually to our clinic for diagnostic evaluation. Since September 2011 whole exome sequencing (WES) is part of the routine diagnostic work-up aimed at the identification of the genetic cause underlying disease[11]. For individuals with unexplained ID, a family-based WES approach is used which allows the identification of *de novo* mutations (DNMs) as well as variants segregating according to other types of inheritance, including recessive mutations and maternally inherited X-linked recessive mutations in males[12]. For this study, we selected all individuals with ID who had family-based WES using the Agilent SureSelect v4 and v5 enrichment kit combined with sequencing on the Illumina HiSeq platform in the time period 2013-2018. This selection yielded a set of 2418 individual probands, including 1040 females and 1378 males across 2387 different families. The level of ID ranged between mild (IQ 50-70) and severe-profound (IQ<30).

Families gave informed consent for both the diagnostic procedure as well as for forthcoming research that could result in the identification of new genes underlying ID by meta-analysis, as presented here.

The exomes of 2418 patient-parent trios were sequenced, using DNA isolated from blood, at the *Beijing Genomics Institute* (BGI) in Copenhagen. Exome capture was performed using Agilent SureSelect v4 and v5 and samples were sequenced on an Illumina HiSeq 4000 instrument with paired-end reads to a median target coverage of 112x. Sequence reads were aligned to the hg19 reference genome using BWA version v0.7.12 and duplicate marking by Picard v1.90. Variants were subsequently called by the GATK HaplotypeCaller (version v3.4-46).

The diagnostic WES process as outlined above only reports (*de novo*) variants that can be linked to the individuals' phenotype. In this study, we systematically collected all DNMs in regions in or close to (200bp) a capture target. DNMs were called as described previously[12]. Briefly, variants called within parental samples were removed from the variants called in the child. For the remaining variants pileups were generated from the alignments of the child and both parents. Based on pileup results variants were then classified into the following categories: "maternal (for identified in the mother only)", "paternal (for identified in the father only)", "low coverage" (for insufficient read depth in either parent), "shared" (for identified in both parents)", and "possibly *de novo*" (for absent in the parents). Variants classified as possibly *de novo* were included in this study.

We applied various quality filters to ensure that only the most reliable calls were included in the study, these are described in the quality control section below.

#### Definition of diagnostic lists

For various analyses in this work, we wanted a list of known developmental disorder genes. In order to define this list, we collected diagnostic lists from each centre in late September 2018 and created sets of "consensus known" and "discordant known" genes.

For the DDD cohort, we used the Developmental Disorders Genotype-Phenotype Database (DDG2P) list, which is a curated list of genes specifically associated with developmental disorders. For every gene on the list, DDG2P provides the level of certainty, consequence of the mutation, and allelic status of variants associated with developmental disorders. We downloaded the DDG2P list on 21 September 2018 from https://decipher.sanger.ac.uk/info/ddg2p. In order to define diagnostic genes that act in a dominant fashion, we selected only genes that were considered "probable", "confirmed", or "both DD and IF" (i.e. high levels of certainty of being a true DD-associated gene) and had an allelic status of "monoallelic", "x-linked dominant", "hemizygous", or "imprinted".

GeneDx maintains a continually curated list of genes, used to define reporting categories for clinical exome and genome testing, which have been definitively or putatively implicated in human Mendelian disease, with modes of inheritance noted for each gene. Starting with the September 2018 curation list, those genes with dominant modes of inheritance and definitive implications in disease were manually reviewed to remove any genes with no association to developmental disorders either because of no phenotypic overlap with the inclusion criteria for this study or because the relevant phenotypes were adult onset.

For the list from RUMC, gene panels for intellectual disability, epilepsy, and craniofacial anomalies/Multiple congenital anomalies were designed by multidisciplinary expert teams consisting of a clinical laboratory geneticist, a molecular geneticist, and a clinical specialist. Each set contained all genes known to be associated with the disease. We used the gene panel version of September 2018 (DGD-2.14). From each of the three gene lists, we selected genes with a reported inheritance of “AD”, “AD,AR”, “AD,IMP”, “AD/AR”, “xl”, “XL”, “XLD”, “XLR,XLD”, or “XLR/XLD”.

After mapping to HGNC IDs and symbols, any gene that was considered diagnostic by all three centres was designated as a “consensus known” gene (n=331). For genes on one or two of the diagnostic lists, we considered them “diagnostic known” genes (n=628).

### Joint quality control of dataset

#### *De novo* mutation filtering

We applied the following filters across all three datasets:

- Removed DNMs outside coding regions
- Removed mutations that fell within known segmental duplication regions as defined by UCSC (http://humanparalogy.gs.washington.edu/build37/data/GRCh37GenomicSuperDup.tab)
- Keep only the most severe DNM in each gene per individual according to the consequence severity from VEP[10]

The following filters were applied specifically to each centre. These were chosen to minimise the number of false positive DNMs. The filters were applied in this specific order.

- DDD
  - Autosomes
    - The minor allele frequency (MAF) < 0.01 across all DDD samples, Exome Aggregation Consortium (ExAC)[13], and 1000 Genomes[14] populations
    - Read depth (RD) of child > 7, mother RD > 5, father RD > 5
    - Fisher exact test on strand bias p-value > $10^{-3}$
    - Remove DNM if any two of the following conditions are met:
      - Both parents had ≥ 1 supporting the alternative allele
      - There is an excess of parental alternative allele within the cohort at the DNMs position. This is defined as p-value <$10^{-3}$ under a one-sided binomial test given an expected site error rate of 0.002
      - There is an excess of alternative alleles within the cohort for DNMs in a gene. This is defined as p-value < $10^{-3}$ under a one-sided binomial test given an expected site error rate of 0.002
    - Filter only applied to indels:
      - Remove the indel if all three conditions are met:
        - Variant allele frequency (VAF) in child < 0.2
        - MAF > 0 for any of the following cohorts: across all DDD samples, ExAC, 1000 Genomes populations
        - Size of indel < 5bp
    - Posterior probability of being a *de novo* mutation (output from DeNovo Gear) > 0.00781 for autosomal DNMs. These thresholds have been

determined through earlier work such that the observed number of synonymous DNMs match the expected number[3,15]

- Filter out mutations in sites with more than one mutation with VAF < 0.3

- X chromosome
  - We ran DeNovoGear as previously described, but with a different set of hard filters to account for the lower coverage in males and to maximise sensitivity and specificity. We examined all candidate DNMs in males and a large subset of those in females manually in IGV[16], and used this to settled on the following set of filters:
    - We removed DNMs in the pseudoautosomal regions.
    - The variant had to be called heterozygous or, for males, hemizygous in the child in the original GATK calls, and called homozygous reference in the parents.
    - We removed variants in segmental duplications.
    - For male probands, we required the following: in the child, alternate allele depth > 2 and RD > 2; in the mother, RD > 5
    - For female probands, we required the following: in the child, alternate allele depth > 2, RD > 7; in the mother, RD > 5; in the father, RD > 1
    - For single nucleotide variants, we required $p > 10^{-3}$ on a Fisher's exact test for strand bias, pooling across trios (ignoring fathers of male probands) where a *de novo* was called at the same site by DeNovoGear.
    - For female probands, we removed indels < 5bp if they had VAF < 0.3 or MAF > 0, since these were vastly over-represented and seemed to be a common error mode.
    - Removed DNMs if any two of the following conditions were met (these conditions were applied separately for males and females):
      - Lowest alternative read count for the parents (or, for males, the mothers) is higher than the maximum allowable given the depth, an error rate of 0.002, and a probability threshold of 0.98 (using the mindepth function in DeNovoFilter)
      - An excess of parental (or, for males, maternal) alternative alleles with a putative *de novo* at that site, defined as p-value < $10^{-3}$ under a one-sided binomial test given an expected error rate of 0.002

        - An excess of parental (or, for males, maternal) alternative alleles with a putative *de novo* in the same gene, defined as p-value > $10^{-3}$ under a one-sided binomial test given an expected error rate of 0.002
    - We set a cut-off for the ppDNM from DeNovoGear to > 0.00085 based on matching the observed number of synonymous DNMs in females to the expected number

- GeneDx
    - RD > 10 for child, mother, and father
    - VAF > 0.15 for child for SNVs and VAF > 0.25 for indels
    - More than 3 reads supporting the alternative allele
    - Genotype Quality (GQ) score > 40
    - Phred-scaled p-value using Fisher's exact test to detect strand bias < 30
    - Log odds of being a true variant versus being false from VQSR > -10 outputted from GATK
    - Any variant with general population frequency above 0.01 was also excluded based on 1000 Genomes and Exome Aggregation Consortium (ExAC) variant population frequency data
    - Filtered out de novo variants called > 4 times in the parental samples in the cohort
    - Filter out DNMs with VAF < 0.3 and VQSLOD < 7 where VQSLOD is the log odds ratio of being a true variant outputted from GATK
    - Filter out *de novo* indels > 100bp
    - Filter out DNMs, not on chromosome X, with a VAF of 1
    - Filter out 5 individuals with more than 10 coding DNMs. These appeared to be due to relatively poor sample quality.
- RUMC
    - GATK Quality score > 450
    - Minimal number of variant reads: 10
    - Minimal number of total reads: 20
    - Minimal percentage of variant reads: 20%
    - Frequency in dbSNP < 0.1%
    - Coverage in parents of at least 10 reads
    - We discarded 15 complex variants after manual inspection in IGV

#### Duplicate samples

We selected a set of 50 common exonic SNPs (only 47 of which could be evaluated across all three centres) and collected genotypes at these SNPs for every sample with a *de novo* mutation found in another individual in the joint set (n=781 DDD, 1307 GeneDx, and 164 RUMC). We then used the gtcheck function from bcftools (https://samtools.github.io/bcftools/bcftools.html) to find discordance between each pair of samples. Pairs with low discordance were manually confirmed, leading to a total of 8 duplicate samples identified. One individual from each duplicate pair was removed from the analyses, leaving 31,058 samples for downstream analyses.

#### Removing variants from siblings

Siblings will sometimes share DNMs that arose as the same mutational event in one of their parental germlines. To avoid double counting these shared DNMs, we identified siblings in each cohort and randomly removed one of each pair of shared variants. In total, we removed 11 DNMs found in siblings.

Additionally, we removed 20 false positive variants identified in novel genes (see below). After filtering we had a total set of 45,221 coding DNMs (**Sup Table 1**). The filtered VAFs across the three datasets are displayed in **Sup Fig 1**.

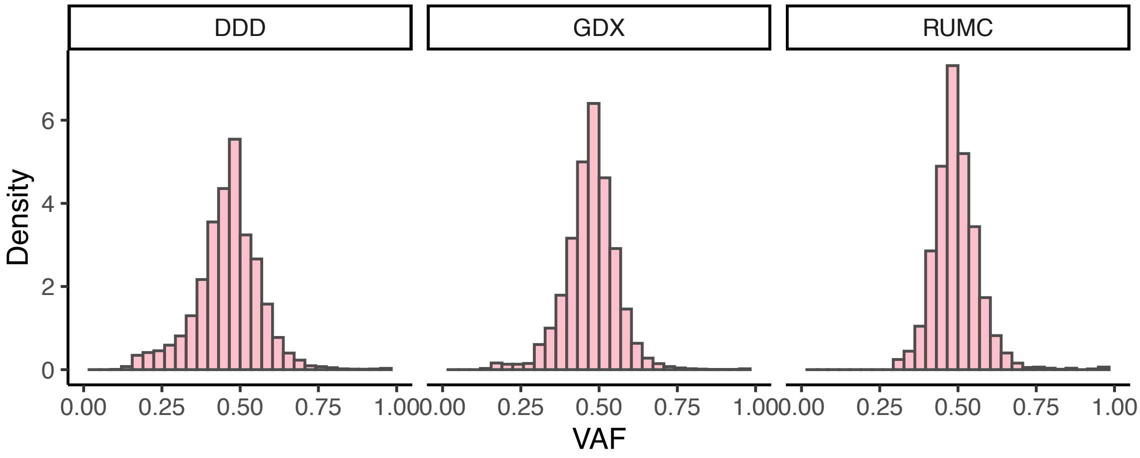


**Supplementary Figure 1**: Variant allele fraction (VAF) for all *de novo* coding mutations, split by centre. Note that mutations with a VAF of ~1 are hemizygous *de novo* mutations in males. DDD = Deciphering Developmental Disorders. RUMC = Radboud University Medical Center.

**Supplementary Table 1**. *De novo* mutations from 31,058 individuals with developmental disorders. For every *de novo* mutation, we provide: proband ID (‘id’), chromosome (‘chrom’), position in GRCh37 (‘pos’), the reference allele (‘ref’), the alternative allele (‘alt’), the VEP consequence of the mutation (‘consequence’), the HGNC symbol (‘symbol’), the centre which sequence the proband (‘study’), the fraction of reads that are from the alternative allele (‘altprop_child’), and the HGNC ID (‘hgnc_id’).

### Comparing datasets

All three cohorts are comprised of individuals with severe developmental disorders. We found that the cohorts had comparable rates of male to female probands (55-57% male cohorts; **Sup Fig 2a**) as well as similar DNM rates (average 1.81-1.96 per individual exome). The DDD study has a significantly higher rate of synonymous DNMs (0.31 *de novo* synonymous DNMs per exome) compared to individuals from GeneDx (GDX; 0.28 per exome; Poisson rate test p = 5.2 x $10^{-7}$) or Radboud University Medical Center (RUMC; 0.28 per exome; Poisson rate test p = 0.0132), which is likely due to differences in *de novo* identification pipelines (**Sup Fig 2b**). Specifically, as described in McRae et al[3] and mentioned above, the DDD study selected a ppDNM (posterior probability of a *de novo* mutation) threshold such that the observed number of synonymous DNMs matched the expected number.

All three cohorts have far more carriers of nonsynonymous DNMs in the consensus genes than expected based on a null mutational model (**Sup Fig 2c**). The specific rate of such carriers differs between cohorts, with GeneDx showing the lowest fraction of such cases, which can be explained by the varying ascertainment between centres.

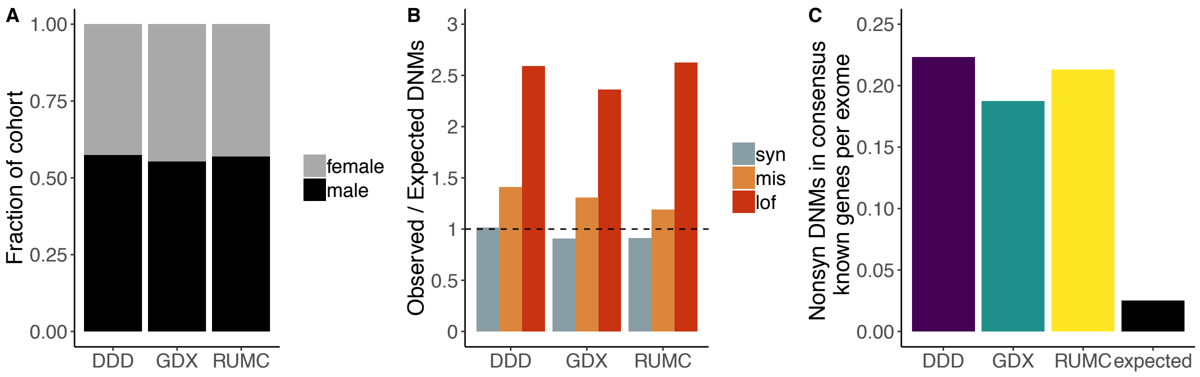


**Supplementary Figure 2**: Comparing cohorts from the three centres. A) Fraction of each cohort that is female (grey) vs male (black). B) Enrichment of observed *de novo* mutations compared to the expected number from a sequence-context based mutational model[17]. C) Rate of nonsynonymous *de novo* mutations (excluding inframe indels) in consensus known genes in each cohort as well as the expected rate based on the aforementioned mutational model. DDD = Deciphering Developmental Disorders. GDX = GeneDx. RUMC = Radboud University Medical Center. Syn = synonymous. Mis = missense. Lof = loss-of-function (including nonsense/stop gained, essential splice site, and frameshift variants).

#### Confirming variants in novel genes

We wanted to confirm all DNMs in the significant genes not previously considered diagnostic by any centre (the “novel” genes), including those genes that were significant in the undiagnosed-only analyses (n = 49 genes). Each centre validated every DNM in these genes either via Sanger sequencing or manual inspection of IGV plots. Across the three centres, only 21 variants appeared to be false positives; these variants were removed from subsequent analyses. The final set of 45,221 *de novo* mutations is included as **Sup Table 1**.

### Enrichment framework

#### DeNovoWEST

DeNovoWEST (*De Novo* Weighted Enrichment Simulation Test) is the testing framework we used to asses gene-wise *de novo* mutation enrichment. Each observed DNM in our dataset was assigned a mutation severity score. This severity score is a proxy for how deleterious we expect the mutation to be. Details of how these were calculated are given below. For each gene we then calculated a gene severity score which is the sum of all mutation severity scores that fall in to that gene.

We used a simulation-based approach to evaluate whether these observed gene severity scores are higher than what we would expect under the null hypothesis of no *de novo* mutation enrichment. To calculate the probability of observing a gene score that is as or more extreme than the one that we observe for this gene (the p-value) we considered the case of observing *k* number of DNMs in the gene where *k* ranged from 0 to 250. This upper limit was chosen as it is far above the number of DNMs we see in any individual gene in our dataset and the probability of observing more than that number of DNMs for our cohort in a single gene is negligible with respect to our significant threshold. The enrichment p-value, pEnrich, was then calculated as the sum of these probabilities as summarized by the following equation where *S* denotes the gene score, *s* is the observed gene score and *K* is the number of DNMs in the gene:

$$P(S \geq s) \approx \sum_{k=0}^{250} P(S \geq s|k)P(K=k)$$

These probabilities were calculated as follows:

- We analytically calculated the probability of observing 0 mutations in this gene and the probability that the severity score of 1 mutation was greater than what we observed. This was calculated based on our sample size and the mutation rate of the gene assuming mutations follow a Poisson distribution.
- To calculate the probability of observing an as or more extreme gene score if we see 2 to 250 DNMs in this gene. We multiplied the probability of observing *k* mutations by the probability of observing a gene score greater or equal to the one we observe. To calculate the latter probability, we simulated the distribution of gene scores as follows:
    - Simulate *k* DNMS across the gene. The probability of mutation was weighted by the trinucleotide sequence context at every base position in that gene.
    - Assign the simulated *de novo* mutations a mutation severity score

- Sum the simulated mutation severity scores to get the simulated gene severity score

#### Determination of Weights

We calculated the weights used in the DeNovoWEST test from observed enrichments across consequence classes, missense constraint information and, for some, CADD score bins (version 1.0)[18]. Enrichments were calculated by dividing the number of observed *de novo* mutations by the number of expected *de novo* mutations across all sites in the exome that fell into a specific stratum. We calculated the number of expected mutations given our sample size and the triplet context mutation rates at the sites[17]. Details for the weight calibration for each consequence class are given below:

- Missense mutations were stratified based on whether or not they fell into a region of missense constraint[19] and into CADD score bins of size 2.5. We fit two LOESS lines on the enrichments for mutations that fall into a region of missense constraint and those that do not.
- Nonsense mutations were stratified based on whether or not they fell into a gene that had a region of missense constraint and into CADD score bins of size 10. We fit two LOESS lines on the enrichments for mutations that fall into a gene with a region of missense constraint and those that do not.
- Synonymous mutations were stratified based on whether or not they fell into a gene that had a region of missense constraint.
- Canonical splice site variants were stratified based on whether or not they fell into a gene that had a region of missense constraint.
- Inframe indels were assigned weights based on the overall enrichment of missense mutations as an appropriate approximation for their deleteriousness. These were stratified by whether or not they fell in a gene with a missense constraint region but not stratified by CADD score bins.
- Frameshift indels were assigned the same weights as nonsense mutations with a CADD score > 40.

These enrichments are depicted in **Sup Fig 3**. The enrichment values or odds ratio (OR) for each stratum were normalised by the level of synonymous enrichment and converted into a positive predictive value (PPV) using the following formula:

$$PPV = \frac{OR - 1}{OR}$$

The synonymous variants were artificially given a PPV of 0.001 as we would expect 1 in 1000 synonymous DNMs in our cohort to be pathogenic. The PPV weights are depicted

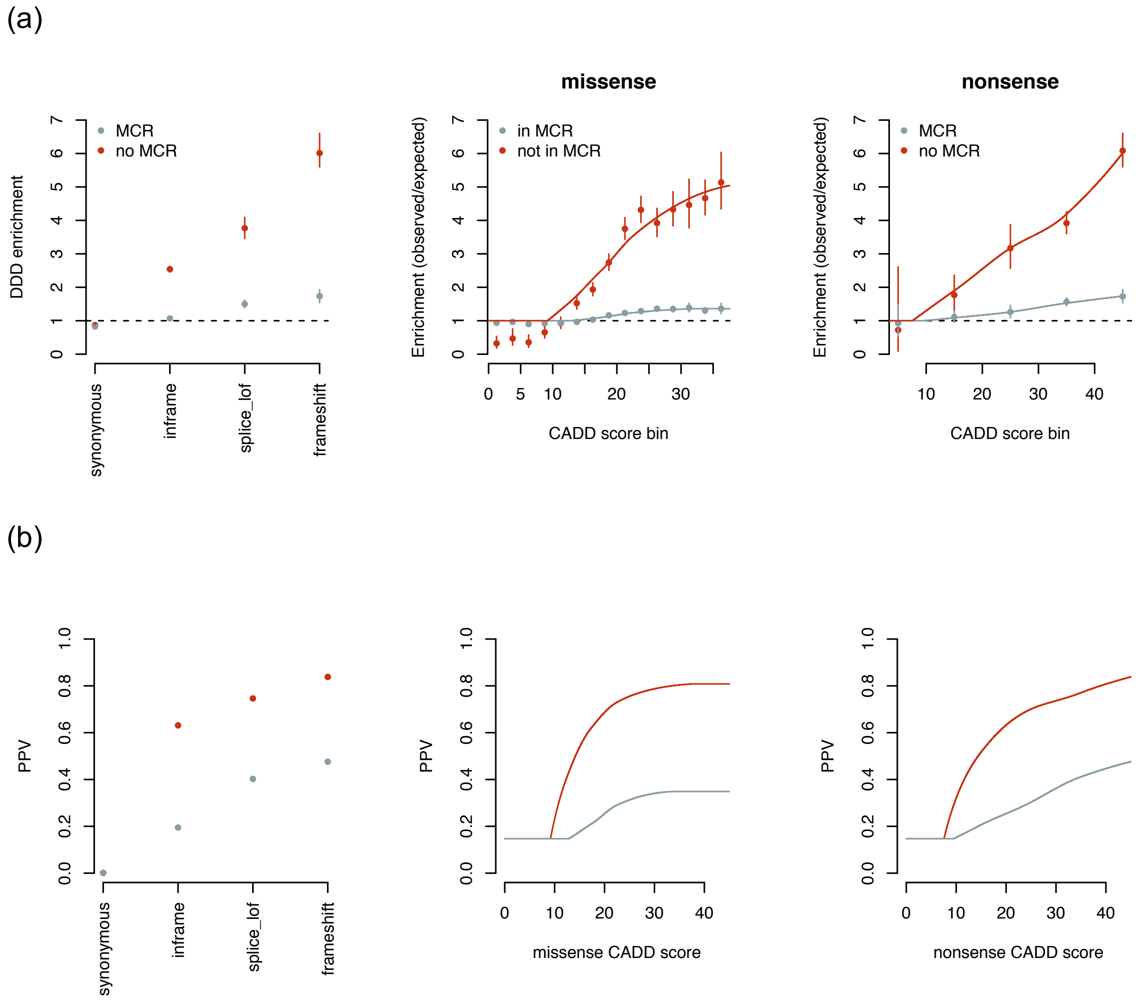


**Supplementary Figure 3: Enrichment of consequence classes and corresponding PPV weights used for DeNovoWEST test.** (a) Depicts the observed enrichment in each consequence class with 95% confidence intervals. The lines fit in the missense and nonsense class are LOESS fits with a restriction not allowing the line to dip below one. (b) Depicts the PPV derived from these observed enrichments. In all these plots points and lines are coloured red if they occurred in a missense constrained region (MCR) (for missense and nonsense variants) or a gene with a region of missense constraint (for the rest of the points). The blue indicates variants in regions or genes without missense constraint.

#### De Novo Clustering Test

We assessed clustering of missense *de novo* mutations within genes to identify genes where DNMs may be acting through dominant negative or activating mechanisms. For this we used DeNovoNear, which has been described previously[3] and is available on GitHub (https://github.com/jeremymcrae/denovonear). We refer to this clustering p-value as pClustering.

#### Combining Tests

We combined the results from the gene enrichment test (pEnrich) and the missense clustering test (pClustering) using Fisher's method. The final p-value was then taken as the minimum of this combined p-value and pEnrich.

#### Independent Hypothesis Weighting Multiple Testing Correction

To correct for multiple testing, we used Independent Hypothesis Weighting (IHW)[20]. IHW assigns weights based on covariates that are informative of the prior probability of the null hypothesis but independent of the p-values. Here we used a weight related to $s_{het}$[21], which is the estimate of selective effect for heterozygous PTVs. Specifically, we used the residuals when we regressed $s_{het}$ on gene length and mutability. This was to ensure that the weight was independent of the p-values. The final set of DeNovoWEST p-values after IHW are listed in **Sup Table 2**. The full framework for DeNovoWEST is depicted in **Sup Fig 4**.

**Supplementary Table 2**. Results of DeNovoWEST. Contains results from analysis on full cohort and on undiagnosed subset along with gene level DNM counts per consequence. Column headers are described within the file.

**Supplementary Table 3**. Novel genes. For each of the 49 novel genes in this analysis, we determined if it had an associated phenotype in OMIM, any publications about an association between mutations in that gene and developmental disorders, and whether it was significant in a study of inherited and *de novo* mutations in autism spectrum disorders[22] ("sig_ASD") or a meta-analysis of *de novo* mutations in individuals with neurodevelopmental disorders[23] ("sig_meta").

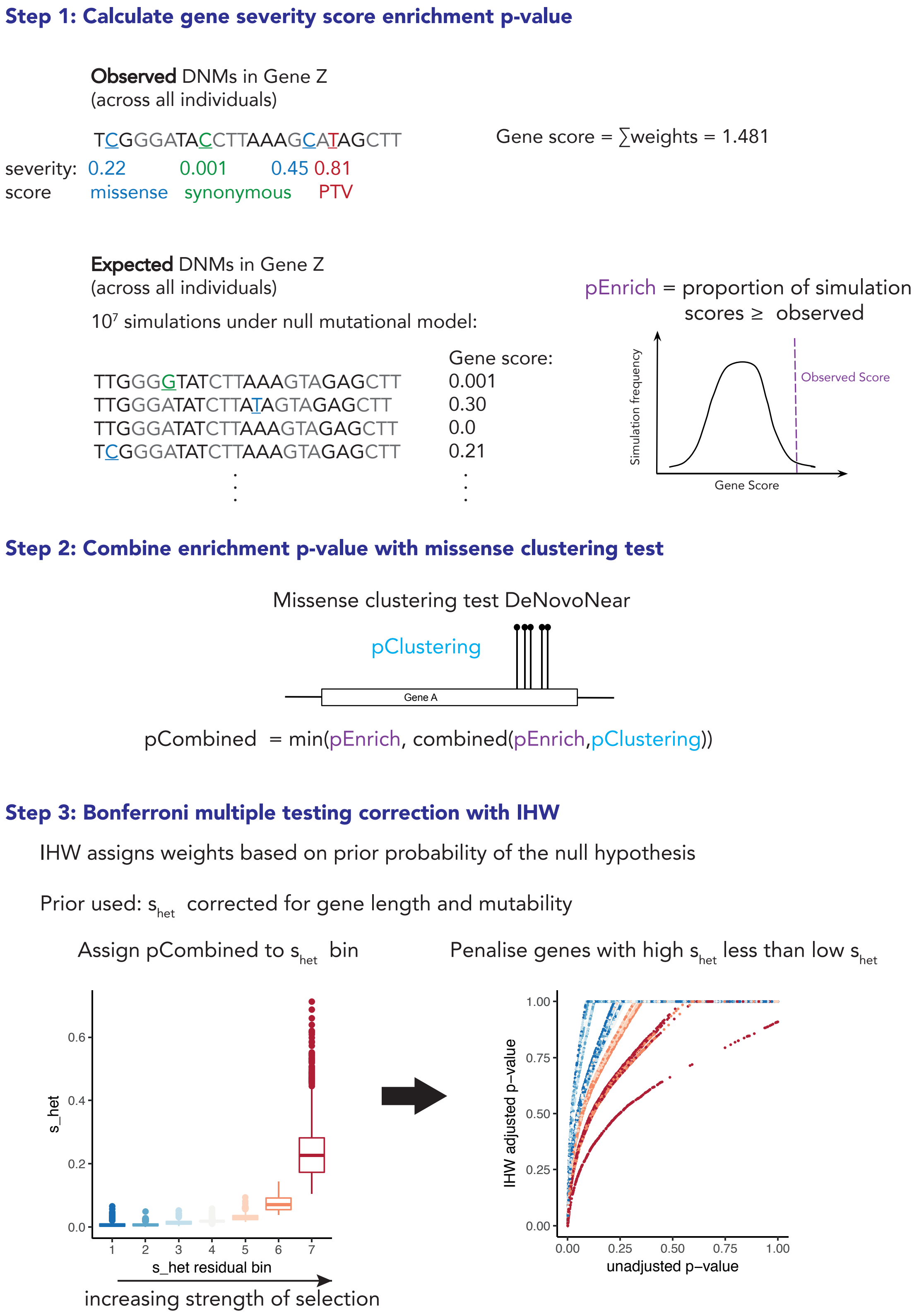


**Supplementary Figure 4: Overview of DeNovoWEST method.**

#### Comparing DeNovoWEST to previous enrichment test

To evaluate our new statistical framework, we ran DeNovoWEST on the DNMs from the ~4k DDD trios from the 2017 publication[3] (not including those from the meta-analysis). We compared the results of the old and new test (**Sup Fig 5**). Using the previous enrichment test, 83 genes were found to be significantly associated with DD while DeNovoWEST identified 96 significant genes. Eighty of these genes overlapped with the previous results, 16 genes were significant in DeNovoWEST results that were not previously and 3 genes that were previously significant were no longer significant. These three genes that we are lost are consensus known genes (*NFIX*, *WWF1A2*, *ITPR1*) that were close to the threshold. For the 16 genes that were gained with DeNovoWEST, 11 were consensus known, 4 were discordant known and 1 was not on any of the diagnostic lists. This demonstrates that we gain substantially more genes than we lose when applying our new enrichment framework.

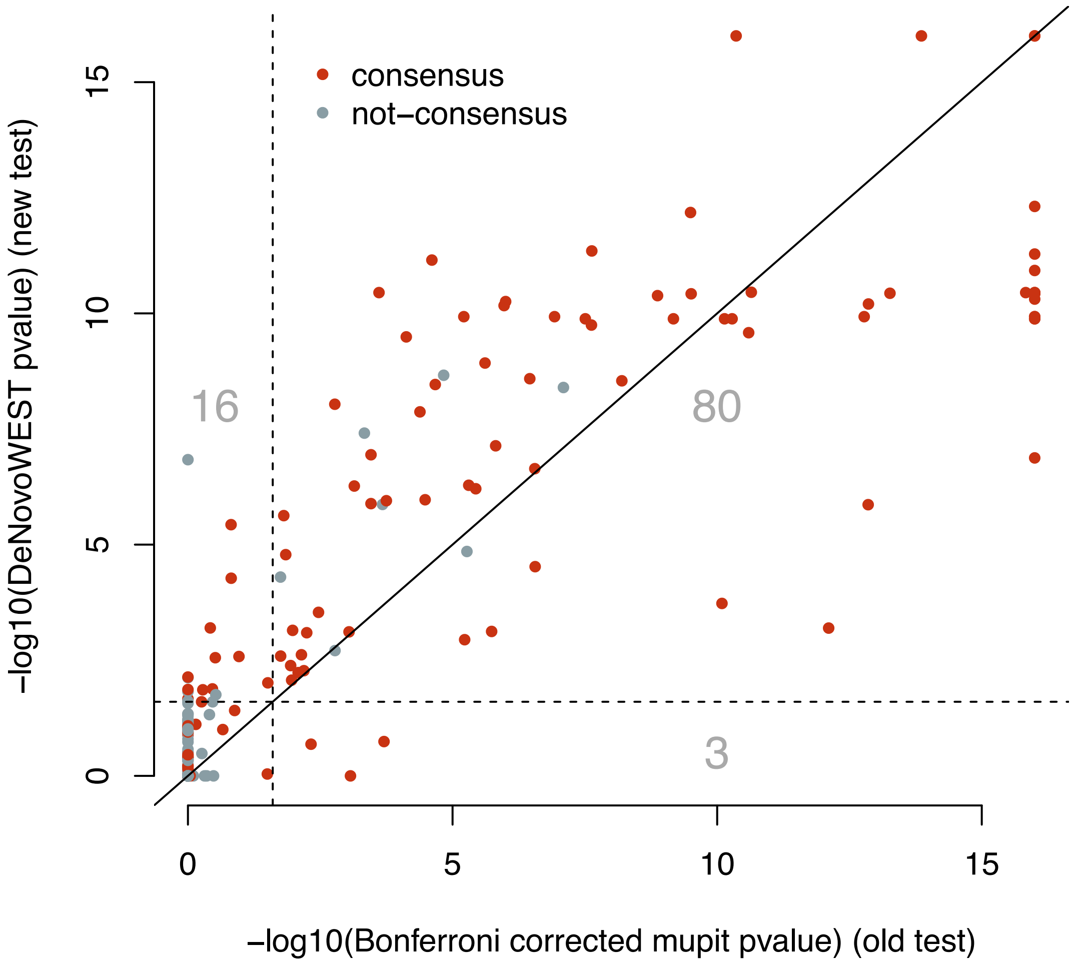


**Supplementary Figure 5:** Figure comparing p-values from published results on 4k DD trios to re-analysis of this data with the new method (DeNovoWEST). Due to constraints on simulation number we do not achieve p-values < $10^{-16}$, therefore the old results are capped at this value for appropriate comparison.

#### Synonymous-only enrichment test

As part of quality control, we repeated the enrichment framework using only synonymous DNMs. While synonymous mutations can be pathogenic, we expect that—as a class—they will not be significantly enriched in any gene. There were 6029 genes with a *de novo* synonymous mutation, but none of these genes was significantly enriched (enrichment $p < 2.66 \times 10^{-6}$, Bonferroni corrected for 18,782 tests; **Sup Fig 6**). Here, we are using the p-value provided after the simulation segment of DeNovoWEST.

Of note, the gene with the highest synonymous enrichment p-value from DeNovoWEST is *KAT6B* (synonymous enrichment $p = 3.1 \times 10^{-5}$), which contains 9 *de novo* synonymous mutations in the joint set. Six of those 9 synonymous variants are the known pathogenic synonymous variant (p.Pro1049Pro) that causes Say-Barber-Biesecker/Young-Simpson syndrome via aberrant splicing of *KAT6B*[24].

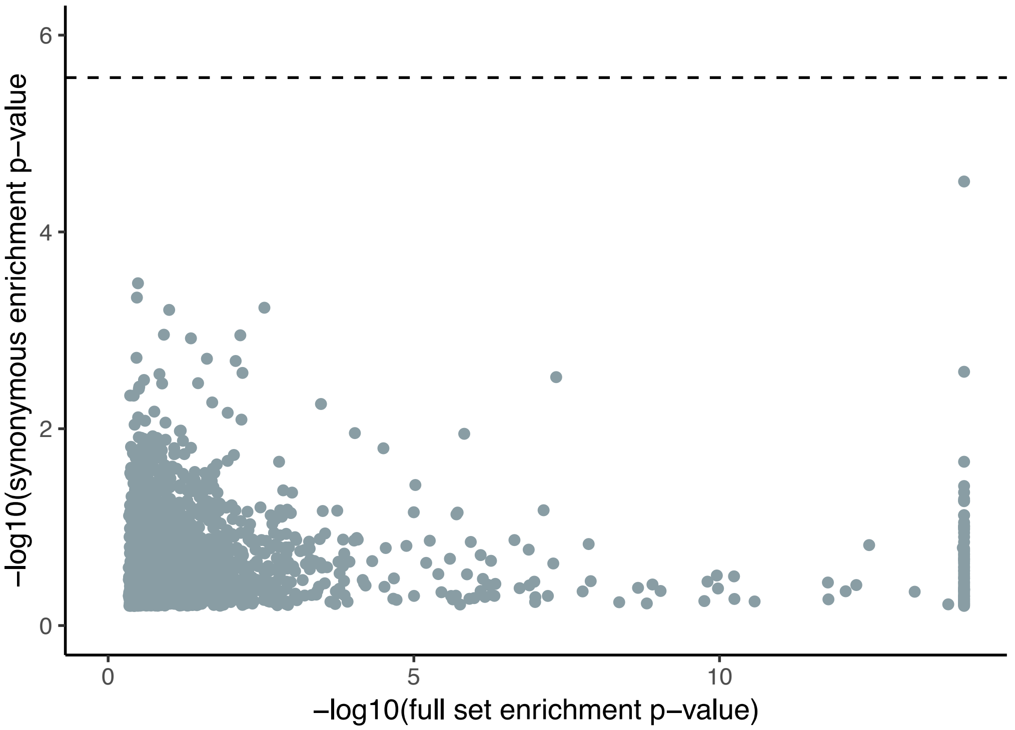


**Supplementary Figure 6.** Comparison of enrichment p-values from the full analysis vs the synonymous-only analysis. Genes with a p-value of 0 have been removed from the plot for clarity.

#### Analysis of undiagnosed cases

We defined diagnosed cases as those carrying a nonsynonymous DNM in one of the 331 consensus known diagnostic genes (n=6211 individuals; 3193 female and 3018 male). We then repeated the enrichment analysis only the remaining 24,847 'undiagnosed' cases with the same weights generated from the analysis of the full set of individuals.

There is a strong correlation between DeNovoWEST p-values in the full dataset compared to those in the undiagnosed-only analysis (**Sup Fig 7**; r = 0.729 for all genes with non-NA DeNovoWEST p-values in both analyses). In total, 118 genes were significant in the undiagnosed-only analysis (DeNovoWEST p < 0.025). Of the significant genes, 69 were discordant known and 49 were novel. As mentioned above, all DNMs in the novel genes were confirmed via Sanger sequencing or IGV inspection.

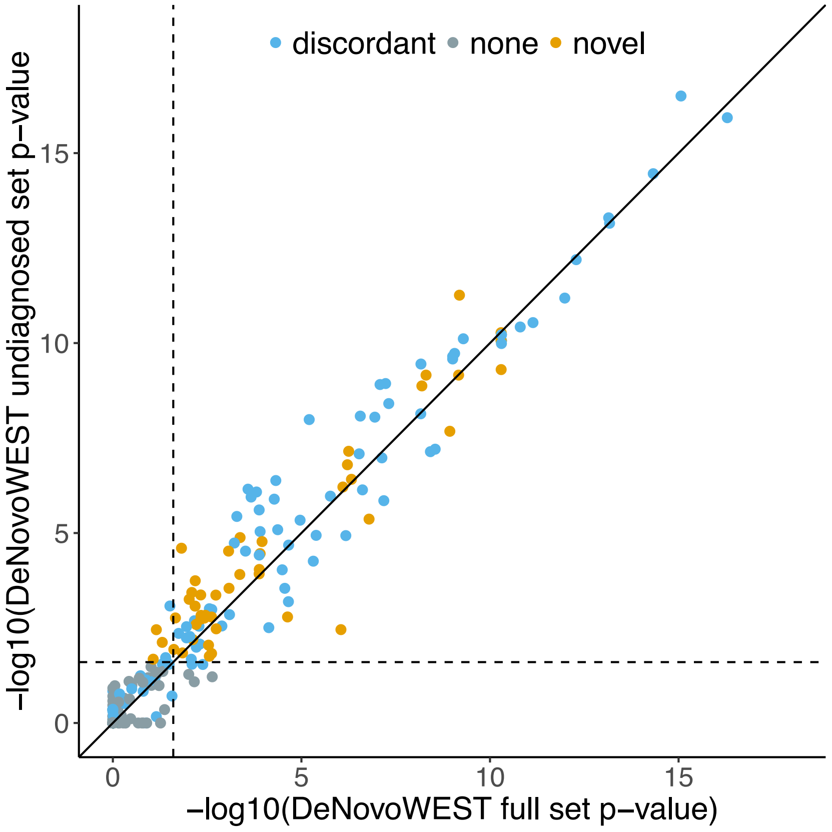


**Supplementary Figure 7**. Comparison between DeNovoWEST p-values for the full analysis vs undiagnosed-only analysis. Note that consensus known genes have been removed since individuals with *de novo* nonsynonymous mutations in these genes were considered diagnosed and removed from the undiagnosed-only analysis. Genes are coloured by their diagnostic list (discordant known = blue; novel = orange; no list / none = grey).

#### Withholding the 10 most mutated genes from weight creation

One potential weakness of our approach is the circularity in defining weights based on mutation enrichments in our cohort and then applying those weights to the same data. Ideally, we would generate weights with each gene held out and then apply weights to the held out gene. However, we hypothesized that doing so would have a minimal impact on the final enrichment results. To test this, we removed the mutations in the 10 most mutated genes (*DDX3X*, *ARID1B*, *ANKRD11*, *KMT2A*, *MECP2*, *DYRK1A*, *SCN2A*, *STXBP1*, *MED13L*, *CREBBP*), regenerated weights, and applied those weights to the full dataset.

We found that the DeNovoWEST p-values for the full analysis vs this analysis were highly correlated ($r = 0.977$ for genes with $p < 1$), including nearly identical p-values for the 10 genes withheld from weight creation. There was only one gene that was significant in the full analysis but not in this analysis (*ASXL2*; $p = 0.0181$ in the full analysis; $p = 0.0318$ in the analysis with the top 10 genes removed from weight calculation). No genes were significant in this analysis but not in the full analysis. These minimal differences support that our approach is robust to this form of circularity.

#### Evaluating potential cryptic splice site variants

While it is known that the terminal exonic G before a splice site can have a strong impact on splicing and be involved in developmental disorders[25,26], there are too few of them as a class to treat as a separate group when determining weights in the enrichment framework. To address these bases and others that may act as cryptic splice sites, we downloaded all publicly available SpliceAI scores[27], which were provided for single nucleotide variants with a maximum delta score ≥ 0.1. We annotated our DNMs with the largest SpliceAI delta score of the four possible scores (acceptor gain, acceptor loss, donor gain, and donor loss). Only 8% (n = 3329/40992) of our single nucleotide DNMs could be annotated with a SpliceAI delta score, but 92% of single nucleotide splice acceptor and donor variants (n = 948/1032) received a score. We selected a threshold of 0.8 to define a set of potentially splice disrupting variants based on three major lines of evidence from Jaganathan, Kyriazopoulou, McRae et al[27]:

1. Exons that are nearly always included in transcripts (high exon inclusion rate) have higher maximum delta scores
2. Higher delta scores were associated with a higher validation rate (but lower sensitivity) when studying GTEx RNA-seq data
3. Variants with higher delta scores (specifically those with a score ≥ 0.8) showed a lower rate of tissue specific usage of reference vs aberrant transcripts

In our data, 1059 DNMs have a delta score ≥ 0.8; the vast majority (83%; n = 874) of these are splice acceptor and donor variants. There were 43 synonymous DNMs with high (≥ 0.8) SpliceAI delta scores that were found in 36 unique genes. The only genes with multiple such synonymous DNMs were *KAT6B* (n = 6), *ARID1B* (n = 2), and *DDX3X* (n = 2), all of which are on the consensus known list and significant in the full analysis. The other 33 genes would gain at most one additional potential PTV. However, one additional PTV would not make any of the non-significant genes significant. As an example, the gene closest to significance was *RAB5C* (simulation enrichment p = 5 x $10^{-5}$; DeNovoWEST p = 0.47). A classic burden test with the synonymous variant considered as a PTV does not survive Bonferroni correction (unadjusted Poisson p = 4 x $10^{-5}$).

Additionally, we extracted the mutational context for all sites with a SpliceAI delta score ≥ 0.8 and used the sample size and triplet context mutation rates[17] to determine the expected numbers of DNMs in our cohort. There is a significant enrichment of DNMs that have a SpliceAI ≥ 0.8 (1.76 fold enriched, Poisson p < $10^{-50}$; **Sup Table 4**), which remains when removing known splice sites (1.55 fold enriched; Poisson p = 2 x $10^{-8}$). The enrichment of synonymous DNMs with high SpliceAI scores is also significant (1.56 fold enrichment; Poisson p = 0.0037;

**Sup Table 4**). The six *KAT6B* synonymous variants mentioned in the previous paragraph (and included in the 43 variants) are a known pathogenic synonymous variant[24]. If those six variants are removed from the enrichment calculations, the p-value is only nominally significant (p = 0.0328, 1.35 fold enriched).

| | Observed DNMs | Expected DNMs | Enrichment | Poisson p-value |
|---|---|---|---|---|
| **All variants** | 1059 | 601.56 | 1.76 | $1 \times 10^{-63}$ |
| **All non-splice variants** | 185 | 119.23 | 1.55 | $2 \times 10^{-8}$ |
| **Synonymous variants** | 43 | 27.48 | 1.56 | 0.0037 |

**Supplementary Table 4**. Comparing observed to expected variants with SpliceAI delta scores ≥ 0.8. The sample size (n = 31,058) and triplet context mutation rates[17] were used to determine the expected number of DNMs for sites with high SpliceAI delta scores (≥ 0.8)[27]. Poisson p-values were computed comparing the observed number of mutations to the expected number. For "All non-splice variants", DNMs with the consequences of "splice_acceptor_variant" and "splice_donor_variant" were removed.

#### Estimating the fraction of cases explained by *de novo* mutations

We previously estimated that the proportion of DDD individuals explained by diagnostic *de novo* coding mutations in known or as-yet-undiscovered genes was between 0.461 and 0.486[15]. For this analysis, we define diagnostic variants as *de novo* nonsynonymous mutations in all consensus known genes and in the significant discordant known and novel genes. Overall, 24.8% of our cohort harbours a DNM that would be considered diagnostic by the previous criteria, with the largest fraction of cases harbouring mutations in the 331 consensus known genes (20% of cases; **Fig 1b**).

##### Splitting by cohort

When evaluating each study separately, we observe 27.1% of DDD probands, 23.3% of GeneDx probands, and 26.4% of RUMC probands harbour a nonsynonymous DNM in a consensus known or significant DD-associated gene. There are no significant differences between cohorts in the rate of individuals who harbour these 'diagnostic' DNMs in novel genes (chi-squared p = 0.6665) or discordant known genes (chi-squared p = 0.0643). Consensus known genes, however, show significantly different rates of diagnostic DNM carriers between cohorts (chi-squared p = $1.28 \times 10^{-10}$). This difference is driven primarily by a lower rate in GeneDx samples: 18.8% in GeneDx vs 22% in DDD (chi-squared p = $6.90 \times 10^{-11}$) and 21.3% in RUMC (chi-squared p = 0.0026). Since the GeneDx cohort is composed entirely of referrals, many patients with suspected genetic disorders in high-prevalence genes are removed from the referral pool by receiving a positive diagnosis via targeted single-gene or panel testing. There is no significant difference in rates between diagnostic DNM carriers in DDD and RUMC for the consensus known genes (chi-squared p = 0.4966). For individuals with multiple damaging DNMs, we gave priority first to DNMs in consensus genes then discordant genes and finally to novel genes.

##### Splitting by sex

When we split the cohorts by the sex of the proband, we find that a significantly higher fraction of females have a nonsynonymous DNM in an autosomal diagnostic gene (as above, all consensus known genes, significant discordant known genes, and novel genes) when compared to males (OR = 1.14, Fisher's p = $1.4 \times 10^{-6}$). This difference is driven by the autosomal consensus known genes, in which a higher fraction of females harbour nonsynonymous DNMs compared to males (OR = 1.17, Fisher's p = $1.17 \times 10^{-7}$; **Sup Fig 8**). We find no significant differences between males and females in the significant discordant genes (autosomal only; OR = 1.05, Fisher's p = 0.448) or novel autosomal genes (OR = 0.89, Fisher's p = 0.2289). Note that these results are only for autosomal genes since we expect varying diagnostic rates on the X chromosome between males and females.

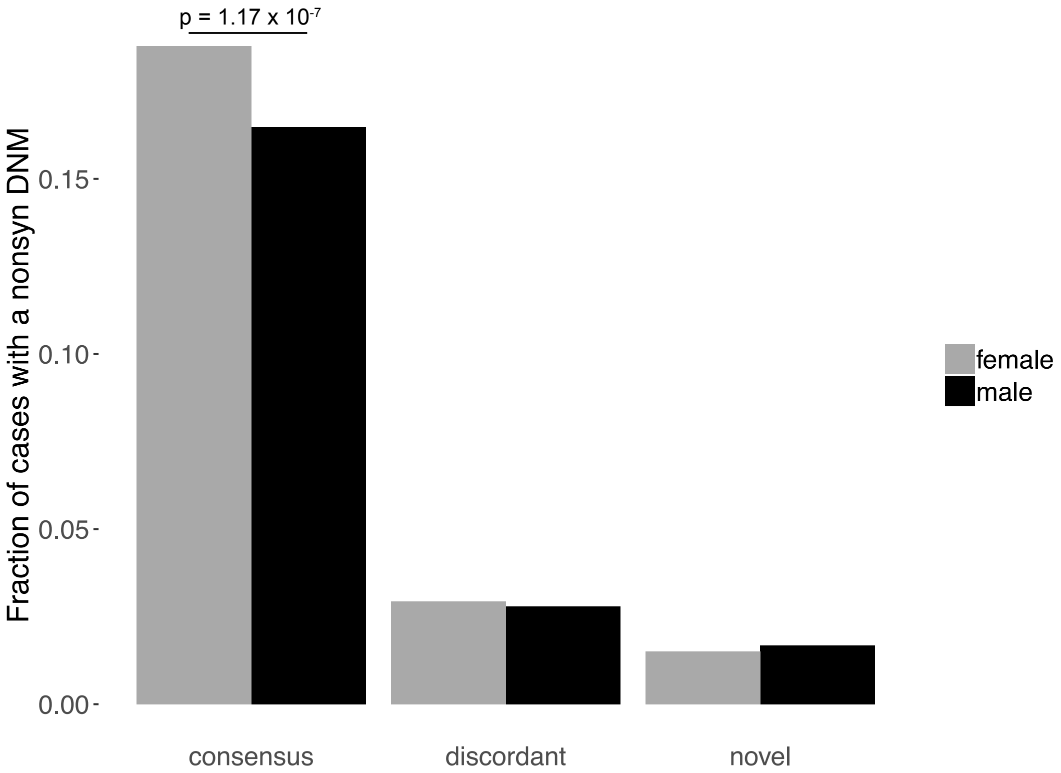


**Supplementary Figure 8:** Fraction of cases with a nonsynonymous DNM, split by sex and diagnostic list. Females are depicted in grey and males in black. Consensus here is all consensus known genes (n = 331). Discordant is only significant discordant known genes (n = 69). Novel is the significant novel genes (n = 49). The only significant difference is for the consensus known genes, where females have a higher rate of nonsynonymous DNMs than males (OR = 1.17, Fisher's p = 1.17 x $10^{-7}$).

When considering all autosomal genes, no significant difference is found between the fraction of female vs male cases who harboured a nonsynonymous DNM (OR = 1.03, Fisher's p = 0.2876). Even when removing individuals with a nonsynonymous DNM in a diagnostic gene (n = 7690), there is no difference in the fraction of female vs male cases carrying a nonsynonymous DNM in autosomal genes (OR = 0.98, Fisher's p = 0.4757). Additionally, we do not observe significantly different proportions of nonsynonymous to all DNMs in autosomal genes in females vs males for either the full set of individuals (p = 0.5346, proportions test) or the subset of individuals without a nonsynonymous DNM in a diagnostic gene (p = 0.6485, proportions test).

### Comparing the new and known genes

#### Phenotypic similarity between new and known genes

Semantic similarity between any two phenotype terms in the Human Phenotype Ontology (HPO) was calculated using the Hybrid Relative Specificity Similarity (HRSS) measure[28]. The information content of each phenotype term was modified by averaging the information content calculated from the HPO phenotype-to-gene annotation file and the information content calculated from the cohort phenotype-to-proband annotations. For this analysis only the most informative phenotypes were used. The set of each proband's phenotypes were pruned so that the final set contained no pairs of phenotype terms where one term was an ancestor of another. The phenotypic similarity between any two probands was defined as the best-match average of HRSS scores for the cross product of the two sets of phenotypes. By definition, both HRSS scores and phenotypic similarity scores range from 0.0 to 1.0.

To compare the phenotypic similarity between the consensus known and novel genes, we computed the phenotypic similarity for each pair of probands who had DNMs (excluding synonymous variants) in the same gene and repeated this for every consensus known and novel gene. There were 134,121 pairwise scores representing comparisons of the probands with DNMs in consensus known and novel genes. For comparison to a null model, we randomly selected 100,000 pairs of probands from a downsampled subset of the entire cohort of 31,058 probands. We found that the random selection of probands is significantly less phenotypically similar than probands with DNMs in novel or consensus known genes (p-value = $2.01 \times 10^{-50}$ and p-value = 0.0, Wilcoxon rank-sum test). The software used to perform the phenotypic similarity calculations is available at https://github.com/GeneDx/phenopy.

#### Functional similarity between new and known genes

To compare the functional similarity between the consensus known and novel genes we looked across various properties that have been known to be important in classifying haploinsufficiency[29]. The details of each variable we have used are detailed below:

- Somatic driver gene: a binary variable of whether the gene is a known somatic driver gene[30].
- Median RPKM fetal brain: the median RPKM in the fetal brain taken from BrainSpan[31].
- Relevant GO term: a binary variable of whether the gene was annotated with one of twenty GO terms that were enriched in consensus DD genes. To select these terms, we annotated all genes with GO terms and looked at the enrichment of each GO term

between consensus DD genes and non-DD genes (genes that are not significant in our analysis and are not on either the discordant or consensus known genes lists). We defined relevant GO terms as the top 20 most enriched terms that appears in at least 20 of the 331 consensus known genes. This was to ensure that we were picking terms that were generalisable to the entire set were not specific to only a few genes. The terms selected are detailed in **Sup Table 5**. At least one of these 20 terms were present in 237 (71%) of consensus genes and 2,874 (16%) of non-DD genes.

- Network distance to consensus known genes: As in Huang et al[29], we created a protein-protein interaction network by integrating information from the Human Protein Reference Database[32], STRING[33], and Reactome[34]. We then calculated the shortest path distance (a measure of proximity) between each gene and consensus known genes.
- Network degree and betweenness: We used MCL[35] (version 14-137; https://micans.org/mcl/) to determine network degree and betweenness (both measures of centrality). Specifically, we used commands mcl query and mcl ctty.
- Promoter GERP and Coding GERP: These were calculated as described in Huang et al.[29]
- pLI: the gnomAD pLI score were used here[36].
- Macaque dN/dS: downloaded from Ensembl.

We compared the mean of these variables across consensus and novel genes compared to non-DD genes (**Figure 2a**) and found that the novel and consensus genes had very similar enrichments across most variables. The biggest difference was found in the network distance to another consensus DD gene (p-value 5.92 x $10^{-5}$, Wilcoxon test).

| GO ID | GO name | Number of non-DD genes with GO term | Number of consensus DD genes with GO term | Enrichment of GO term in consensus vs non-DD genes |
|---|---|---|---|---|
| GO:0001501 | skeletal system development | 120 | 30 | 13.857251 |
| GO:0007507 | heart development | 162 | 32 | 10.948939 |
| GO:0008543 | fibroblast growth factor receptor signalling pathway | 139 | 23 | 9.171706 |
| GO:0001701 | in utero embryonic development | 219 | 35 | 8.858516 |
| GO:0044212 | transcription regulatory region DNA binding | 169 | 27 | 8.855521 |
| GO:0010628 | positive regulation of gene expression | 156 | 23 | 8.172225 |
| GO:0007411 | axon guidance | 297 | 42 | 7.838445 |
| GO:0003682 | chromatin binding | 333 | 46 | 7.656859 |
| GO:0043234 | protein complex | 324 | 41 | 7.014164 |
| GO:0000122 | negative regulation of transcription from RNA polymerase II promoter | 543 | 68 | 6.941385 |
| GO:0007268 | synaptic transmission | 362 | 42 | 6.430989 |
| GO:0045893 | positive regulation of transcription, DNA-dependent | 519 | 59 | 6.301178 |
| GO:0007399 | nervous system development | 286 | 32 | 6.201846 |
| GO:0045944 | positive regulation of transcription from RNA polymerase II promoter | 718 | 78 | 6.021535 |
| GO:0007420 | brain development | 203 | 22 | 6.007084 |
| GO:0005667 | transcription factor complex | 223 | 24 | 5.965453 |
| GO:0048011 | neurotrophin TRK receptor signalling pathway | 256 | 27 | 5.846028 |
| GO:0006184 | GTP catabolic process | 229 | 22 | 5.325057 |
| GO:0008134 | transcription factor binding | 284 | 27 | 5.269659 |
| GO:0003924 | GTPase activity | 276 | 26 | 5.221573 |

**Supplementary Table 5**: Table showing the GO terms selected as being relevant to consensus DD genes.

#### Evaluating potentially smaller mutational targets in the novel genes

We hypothesize that the novel genes may have a smaller mutational target than the previously established genes. In the following set of analyses, we only consider significant consensus known genes (n = 181) and compare them to the significant discordant and novel genes (n = 118).

Many of the known developmental disorder genes act via haploinsufficiency (e.g. > 50% of the monoallelic DDG2P list have "loss of function" as their listed mutation consequence) and could therefore be caused by almost all true loss-of-function variants within a gene. Disorders due to gain-of-function mutations, however, would be caused by a much smaller number of mutations within a gene and would therefore require large sample sizes to be significantly burdened by DNMs. To investigate if the new genes (here both the discordant and novel) were more likely to act via a gain-of-function mechanism, we compared the ratio of missense to PTV DNMs in the gene sets and found that the consensus known genes have a significantly lower missense:PTV ratio than the discordant and novel genes (ratio 1.17 vs 2.04; chi-squared $p = 3.25 \times 10^{-21}$).

Alternatively, a smaller mutational target could be due to a shorter gene length. We would expect this to be most obvious for genes predicted to act via haploinsufficiency. To define such likely haploinsufficient genes, we subsetted the 299 genes to 159 with significant PTV enrichment ($p < 1 \times 10^{-6}$). We, however, do not find significant differences in gene length or PTV mutability between the consensus known genes compared to discordant and novel genes (Wilcox p = 0.9647 and 0.8958, respectively).

#### Comparison of constraint metrics

Additionally, we compared various constraint metrics between the significant consensus known genes and the significant discordant and novel genes. To correct for mutability, for each of these metrics, we assigned missing genes the median value. We then ran a regression of the probability of mutation on the constraint metric and used the residuals in Wilcoxon rank sum tests. Using these residuals therefore corrects for systematic differences in length in between gene sets. When correcting for mutability in this way, we find no significant differences between the two gene sets for $s_{het}$[21] (Wilcox p = 0.9733), pLI[36] (Wilcox p = 0.9831), or LOEUF (Wilcox p = 0.9047). We also find no significant difference between the known and novel genes for the rate of observed/expected missense variants in gnomAD (oe_mis; Wilcox p = 0.7359).

#### Clustering of PTVs at the end of transcripts

We also wanted to search for clustering of *de novo* PTVs specifically at the terminal end of genes since transcripts with such PTVs can sometimes escape from nonsense mediated decay (NMD). To test this, we selected all genes with > 2 *de novo* PTVs in our cohort and determined the probability of a PTV mutation in regions likely to escape from NMD (the last exon and 50bp into the penultimate exon[37]). We then used a Poisson test to compare the rate of PTVs in the NMD escape region to the rate in the rest of the gene. With a Bonferroni corrected threshold of $1 \times 10^{-6}$, this analysis identified two significant genes: *UBE3A* (consensus known gene; Poisson test $p = 1.96 \times 10^{-9}$) and *MN1* (novel gene; Poisson test $p = 1.58 \times 10^{-7}$; **Sup Fig 9c**).

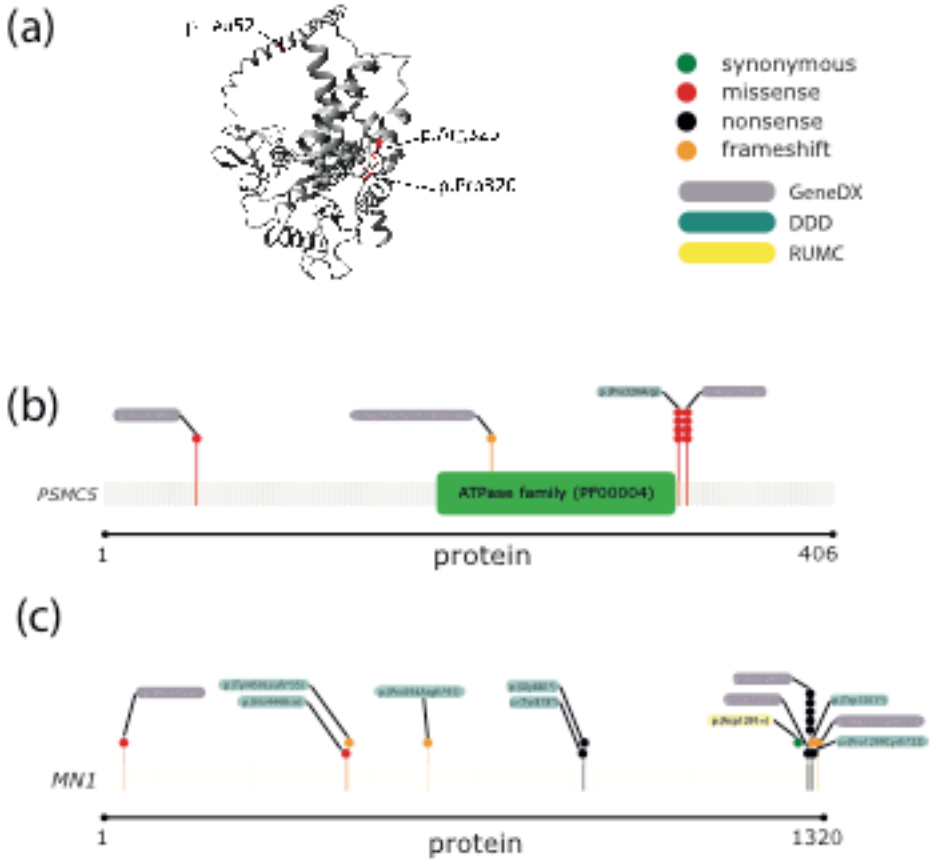


**Supplementary Figure 9**: Models of two novel genes not acting via haploinsufficiency (a) 3D structure (PDB ID: 5GJQ) of novel gene *PSMC5*, highlighting the clustered missense mutations. Modeled and visualized using YASARA & WHAT IF Twinset[38]. Gene models of novel genes *PSMC5* (b) and *MN1* (c). The *de novo* mutations are depicted on each gene model coloured by the type of mutation (synonymous in green, missense in red, nonsense in black, and frameshift in orange). The mutation labels are coloured by the cohort in which the mutation was found (purple for GeneDx, green for DDD, and yellow for RUMC).

### Protein domain analysis

#### Annotation of transcripts, protein positions, and protein domains to genomic coordinates

We used a local version of the MetaDome web server[39] (https://stuart.radboudumc.nl/metadome) to query genomic positions of our variants and obtain corresponding GENCODE[40] transcripts, protein positions in UniProtKB/Swiss-Prot databank entries for the human species[41], and Pfam-A v30.0 protein domains[42].

In order to avoid different variant effect interpretations because of different exon composition and/or reading frames in alternative transcripts, we used a single transcript per gene for the protein domain analysis. A GENCODE transcript was chosen for each gene based on the following criteria in order:

1. The transcript corresponds to a human canonical or isoform entry in Swiss-Prot
2. This transcript contains all (or most) of the *de novo* mutations for the corresponding gene
3. The transcript translates to the longest protein sequence length
4. If multiple transcripts remain for a gene, one of these is selected

All variant consequences were reannotated in context of the chosen transcript. After this selection of transcripts 92.7% of missense, 94.35% of synonymous, and 89.94% of stop-gained variants remained.

#### Counting variants by type and correcting for region-specific bias

The protein domain region coverage (i.e. percentage of a protein that is a protein domain) over all genes (n = 11,983) in the joint *de novo* set is 42.71%. There are on average 2.4 protein domains in each of these genes. Of the 37,089 SNVs in the entire dataset 17,902 (48.27%) are located in a protein domain, and of the variants located in a protein domain, 73.25% are missense, 21.57% are synonymous, and 5.17% are nonsense.

We tested whether specific variant types were more likely to be found within protein domain regions of the cDNA of a gene versus the non-protein domain parts. In this analysis we only focussed on single nucleotide variants in the protein coding part of the gene, and therefore only considered missense, synonymous and stop-gained variants. We counted how often a variant type occurred and in which part of the gene (i.e. protein domain or the remaining gene area). Counts were corrected for the percentage of protein domain region coverage in a gene and for the nucleotide sequence context based on the codon table.

To correct for the protein domain region coverage in a gene we define $P(domain)$ as the percentage of domain region and $P(\neg\ domain)$ as the percentage of the remaining gene region. For the sequence context we extract the nucleotide sequence from the protein domain parts of the gene and use the codon table to compute the probability per codon on variant type $var \in \{'missense', 'synonymous', 'stopgained'\}$. We define the corresponding probability of a variant type in the protein domain region as $P(var_{dom})$ and the probability of a variant in the remaining gene region as $P(var_{\neg dom})$.

The probability of a specific variant type to be found within a protein domain region can then be presented as $P(var_{dom}|\ domain) = P(var_{dom}) \cdot P(domain)$, and $P(var_{\neg dom}|\ \neg domain) = P(var_{\neg dom}) \cdot P(\neg domain)$ for variants in the remaining region of the gene. In general, $P(var_{dom}|\ domain\ ) + P(var_{\neg dom}|\ domain) \neq 1.0$. To use these probabilities as corrections we need to normalize in the following way: $P(var_{dom}|\ domain)^{norm} = \frac{P(var_{dom}|\ domain)}{P(var_{dom}|\ domain)+P(var_{\neg dom}|\ \neg domain)}$ and $P(var_{\neg dom}|\ \neg domain)^{norm} = \frac{P(var_{\neg dom}|\ \neg domain)}{P(var_{dom}|\ domain)+P(var_{\neg dom}|\ \neg domain)}$. The variant counts are then subsequently corrected by counting the variants located in domain regions and multiplying this with $\frac{1.0}{P(var_{dom}|\ domain)^{norm}}$ and for variants found in the remaining gene regions with $\frac{1.0}{P(var_{\neg dom}|\ \neg domain)^{norm}}$ .

| **(a)** | **In domain region** | **In remaining gene region** | **Percentage in domain region** |
|---|---|---|---|
| **Missense (corrected)** | 28,982.13 | 23,799.43 | 54.91 % |
| **Synonymous (corrected)** | 8,358.50 | 8,157.82 | 50.61 % |
| **Stop gained (corrected)** | 2,517.72 | 2,334.40 | 51.89 % |

| **(b)** | **In domain region** | **In remaining gene region** | **Percentage in domain region** |
|---|---|---|---|
| **Missense (corrected)** | 5,797.27 | 2,101.23 | 73.40 % |
| **Synonymous (corrected)** | 299.06 | 283.28 | 51.36 % |
| **Stop gained (corrected)** | 1,192.81 | 943.69 | 55.83 % |

| **(c)** | **In domain region** | **In remaining gene region** | **Percentage in domain region** |
|---|---|---|---|
| **Missense (corrected)** | 432.01 | 230.93 | 65.17 % |
| **Synonymous (corrected)** | 48.12 | 57.28 | 45.65 % |
| **Stop gained (corrected)** | 74.16 | 69.05 | 51.78 % |

**Supplementary Table 6.** (a) Contingency table for all *de novo* single nucleotide variants after correction. G-test: p-value = $3.92 \times 10^{-22}$ (Bonferroni correction: N = 3), test statistic = 100.78, degrees of freedom = 2. Counts of variants are corrected for relative domain fraction of a gene and codon composition. (b) Contingency table for *de novo* single nucleotide variants after correction observed in the consensus known genes G-test: p-value = 1.57 x $10^{-68}$ (Bonferroni correction: N=3), test statistic = 314.45, degrees of freedom = 2. Counts of variants are corrected for relative domain fraction of a gene and codon composition. (c) Contingency table for *de novo* single nucleotide variants after correction observed in the novel genes. G-test: p-value = 1.35 x $10^{-4}$ (Bonferroni correction: N=3), test statistic = 20.02, degrees of freedom = 2. Counts of variants are corrected for relative domain fraction of a gene and codon composition.

#### Computing variant enrichment in protein domains

To determine whether domain families are enriched with variants of a specific type (i.e. $var \in \{'missense', 'synonymous', 'stopgained'\}$), we aggregated the variant counts of each type over the occurrences of that domain family: $obs\_n_{var}(x)$, where$x$represents the domain family. We compare this to the number of expected variants per domain family: $exp\_n_{var}(x)$. We then contrast the observed and expected variant counts in a contingency table to calculate a potential enrichment or depletion via a G-test. Bonferroni correction is applied based on the number of domain families present in the enrichment test set. The contingency table for domain family $x$:

| | **Variants in domain family** | **Variants in other domains** |
|---|---|---|
| Observed variants | $obs\_n_{var}(x)$ | $\sum_{y,y\neq x} obs\_n_{var}(y)$ |
| Expected variants | $exp\_n_{var}(x)$ | $\sum_{y,y\neq x} exp\_n_{var}(y)$ |

The expected number of variants$exp_n_{var}(x)$for a protein domain family is based on a redistribution of$obs_n_{var}(x)$over the entire set of domain families. This redistribution is corrected for any potential bias that could affect the number of variants. Specifically, we correct for the total sequence context of domain regions, the total number of occurrences of a domain family, and the mutation rate of protein regions containing the domain family of interest.

We correct for variant type preference based on the sequence context by extracting the nucleotide sequence of each protein domain region belonging to a family$x$and use the codon table to count the number of possible SNVs that would result into the variant of type $var$: $possible_{var}(x)$. The correction for family $x$ is comparing the total possible SNVs of type $var$ for family $x$ to the total of the entire set of domain families: $correction\_possible_{var}(x) = possible_{var}(x) / \sum_y possible_{var}(y)$.

The number of protein domain occurrences is another aspect we correct for. The number of occurrences of family $x$ is given by$occurrences_x$ and is compared to the total number of protein domain occurrences: $correction\_occurrences(x) = occurrences_x / \sum_y \; occurrences_y$.

The mutation rate of the protein in which a protein domain occurs might be biased. To correct for this we count the total number of mutations found in all the proteins that contain a domain of family $x$ and define that as $prot\_obs\_n_{var}(x)$, then we divide that by the total protein sequence length of all of these proteins $length_prot(x)$to obtain the mutation rate: $mutrate\_prot_{var}(x) = prot\_obs\_n_{var}(x) \; / \; length\_prot(x)$. To correct for the mutation rate we compare it to the total mutation rate for all domains and their corresponding proteins: $correction\_mutrate_{var}(x) = mutrate\_prot_{var}(x) \; / \; \sum_y \; mutrate\_prot_{var}(y)$

An example of the input for assessing domain family enrichment. In this example there are three domain families named A, B, and C:

| Domain family | $possible_{var}(x)$ | $occurrences_x$ | $mutrate\_prot_{var}(x)$ | $obs\_n_{var}(x)$ |
|---|---|---|---|---|
| A | 170 | 3 | 0.3 | 7 |
| B | 160 | 2 | 0.1 | 2 |
| C | 30 | 1 | 0.05 | 10 |
| Total | 360 | 6 | 0.45 | 19 |

The correction variables are computed to calculate the expected number of variants for assessing domain family enrichment:

| Domain family | $correction\_possible_{var}(x)$ | $correction\_occurrences(x$ | $correction\_mutrate_{var}(x)$ |
|---|---|---|---|
| A | 170/360 | 3/6 | 0.3/0.45 |
| B | 160/360 | 2/6 | 0.1/0.45 |
| C | 30/360 | 1/6 | 0.05/0.45 |
| Total | 1.0 | 1.0 | 1.0 |

The expected number of variants is a redistribution of the total number of variants found over all domain families and this is multiplied by each of the correction terms for domain family x. Each of the correction terms are weighed equally and $exp\_n_{var}(x)$ is thus calculated as:

$exp\_n_{var}(x) =$

$$\sum_y obs\_n_{var}(y) \cdot \frac{1}{3} (correction\_possible_{var}(x) + correction\_occurrences(x) + correction\_mutrate_{var}(x))$$

For our example the expected variants are:

| Domain family | $exp\_n_{var}(x)$ |
|---|---|
| A | $19 \cdot \frac{1}{3} \cdot (170/360 + 3/6 + 0.3/0.45) = 10.38$ |
| B | $19 \cdot \frac{1}{3} \cdot (160/360 + 2/6 + 0.1/0.45) = 6.33$ |
| C | $19 \cdot \frac{1}{3} \cdot (30/360 + 1/6 + 0.05/0.45) = 2.29$ |
| Total | 19 |

We assessed if specific domain families are enriched for variant types via this enrichment test for the domains present in the consensus diagnostic genes and these results can be found in **Supplementary Table 7**. We find five domain families enriched with DNM missense variants and one depleted with DNM missense variants. The molecular functions and disease associations due to missense variants of these missense enriched domain families can be found in **Supplementary Table 8**. We did not find any enrichment or depletion when this test was performed for synonymous or stop-gained DNM variants.

| **Domain families** | **Depleted/ Enriched** | **DNM Observed** | **DNM Expected** | **Domain instances** | **Genes** | **Corrected P-value** |
|---|---|---|---|---|---|---|
| Ion transport protein (PF00520) | Enriched | 298 | ~173 | 29 | 15 | 3.89e-07 |
| Ligand-gated ion channel (PF00060) | Enriched | 80 | ~21 | 3 | 3 | 6.71e-07 |
| Protein kinase domain (PF00069) | Enriched | 131 | ~77 | 12 | 11 | 4.44e-02 |
| Protein phosphatase 2A regulatory B subunit (B56 family) (PF01603) | Enriched | 43 | ~12 | 1 | 1 | 1.06e-02 |

| | | | | | | |
|---|---|---|---|---|---|---|
| Sec1 family (PF00995) | Enriched | 47 | ~14 | 1 | 1 | 7.98e-03 |
| WD domain, G-beta repeat (PF00400) | Depleted | 37 | ~81 | 16 | 5 | 1.03e-02 |

**Supplementary Table 7.** Significant missense variant enrichment of domain families in the consensus diagnostic genes (N total domain families = 243, used for Bonferroni correction on the P-value). There is no significant enrichment for synonymous variants.

| Domain | Molecular function | Associated diseases (due to missense) |
|---|---|---|
| Ligand-gated ion channel (PF00060) | Occur as ionotropic glutamate receptors (iGluRs) and plays a critical role in synaptic communication and plasticity (Bortolotto 1999[43], Croset 2010[44]) | (neurological) channelopathies (Catterall 2008[45]) |
| Ion transport protein (PF00520) | Neuronal excitability and synaptic transmission through the central and peripheral nervous systems (Spillane 2016[46]). | (neurological) channelopathies (Spillane 2016[46], Catterall 2008[45]) |
| Protein phosphatase 2A regulatory B subunit (B56 family) (PF01603) | Regulatory function involved in multiple aspects of cell growth and metabolism (McCright and Virshup 1995[47]). | Intellectual disability (ID), Autism, Hypotonia, Macrocephaly (Shang 2016[48]) |
| Protein kinase domain (PF00069) | Regulatory domain that functions as an on/off switch for many cellular processes (Manning 2002[49]) | Associated with many different diseases, including developmental ID (Hamilton 2018[50]) |
| Sec1 family (PF00995) | Neurotransmitter release by exocytosis (Halachmi and Lev 1996[51]). Whereas their interaction with syntaxin molecules represent a negative regulatory step in secretion (Bracher 2000[52]). | Epileptic Encephalopathy (Saitsu 2008[53]) |

**Supplementary Table 8.** Molecular functions and disease associations of the protein domains that are significantly enriched with missense variants.

### Recurrent Mutations

We identified 773 recurrent mutations in our combined dataset (736 single nucleotide variants and 37 indels). The most recurrently mutated sites are listed in **Table 1**.

##### Germline selection genes

We found 39 mutations that fell in 12 known germline selection genes (*FGFR2*, *FGFR3*, *PTPN11*, *HRAS*, *KRAS*, *RET*, *BRAF*, *CBL*, *MAP2K1*, *MAP2K2*, *RAF1*, *SOS1*)[54,55]. To determine if the observed mutations in germline selection genes are known to be activating, we first confirmed that these were all genes known to be acting through gain-of-function mechanisms. All of these genes are known monoallelic DD-associated genes annotated as having activating mutation consequences according to DDG2P[4]. We then confirmed that all these recurrent mutations are listed as pathogenic or likely pathogenic variants in ClinVar[56].

### Overlap with somatic driver genes and mutations

We downloaded a list of 369 somatic driver genes identified in Martincorena et al[30], which we then mapped to their HGNC identifier for comparison with genes in our analysis. There are 73 genes shared between the somatic driver gene list and the 299 significant genes in our full analysis (51 consensus known, 16 discordant known, and 6 novel). This overlap between the two lists is significant when correcting for $s_{het}$[21] (logistic regression, $p = 1.7 \times 10^{-46}$).

To determine the enrichment of DNMs in the somatic driver genes, we used the previously-mentioned sequence context-based mutational model to predict the expected number DNMs in the gene set[13,17]. The somatic driver genes are significantly enriched for nonsynonymous DNMs compared to expectation (missense and PTVs; Poisson $p < 10^{-50}$ for both categories; **Sup Table 9A**). This enrichment remains when removing the 73 somatic driver genes that are significant in the *de novo* analysis (Poisson $p = 7.1 \times 10^{-20}$ for missense and $7.0 \times 10^{-18}$ for PTVs; **Sup Table 9B**). Overall, 9.4% of cases carry a nonsynonymous DNM in a somatic driver gene; in 6.8% of cases, the somatic driver gene impacted is also significant in the *de novo* analyses.

A) All somatic driver genes

| | Observed DNMs | Expected DNMs | Enrichment | Poisson p-value |
|---|---|---|---|---|
| **Synonymous** | 286 | 304.77 | 0.938 | 0.1474 |
| **Missense** | 1810 | 686.17 | 2.638 | $p < 10^{-50}$ |
| **Protein-truncating variant** | 1114 | 94.84 | 11.746 | $p < 10^{-50}$ |

B) Somatic driver genes not significant in the *de novo* analysis

| | Observed DNMs | Expected DNMs | Enrichment | Poisson p-value |
|---|---|---|---|---|
| **Synonymous** | 200 | 215.00 | 0.930 | 0.1614 |
| **Missense** | 690 | 478.48 | 1.442 | $7.1 \times 10^{-20}$ |
| **Protein-truncating variant** | 146 | 65.38 | 2.233 | $7.0 \times 10^{-18}$ |

**Supplementary Table 9**. Enrichment of *de novo* mutations in somatic driver genes. Using a sequence-context based mutational model[13,17] and the size of the cohort, we predicted the expected number of *de novo* mutations (DNMs) by mutational consequence in 369 somatic driver genes[30] (a) and the somatic driver genes that were not significant in the *de novo* analysis (b). These expectations were compared to the observed number of DNMs with a Poisson test.

We took a list of recurrently mutated sites (≥ 3 patients) in the MC3 calls from The Cancer Genome Atlas (TCGA), which included 10,224 cancer patients across 30 cancer types[57]. Overall, we find 117 DNMs at 76 unique sites in our cohort that are recurrently mutated in the TCGA data in at least three patients. To determine enrichment at these recurrently mutated sites in TCGA, we focused on single nucleotide variants only. We extracted the local sequence context and used triplet context mutation rates[17] and the sample size to determine the expected numbers of DNMs in our cohort at these recurrently mutated single nucleotide sites. We performed this analysis while varying the number of patients in TCGA in which the mutation was found (from three or more patients to 10 or more patients). For example, there are 112 DNMs at 71 unique sites in our cohort that are recurrently mutated in the TCGA data in at least three patients (Poisson $p < 10^{-50}$; **Sup Fig 10**). If we were to use a recurrence threshold of 10 in TCGA, however, we find 28 DNMs at 12 unique sites (Poisson $p = 1.2 \times 10^{-39}$). As the recurrence threshold in TCGA is increased, an increasingly large proportion of the DNMs are recurrently mutated in our cohort.

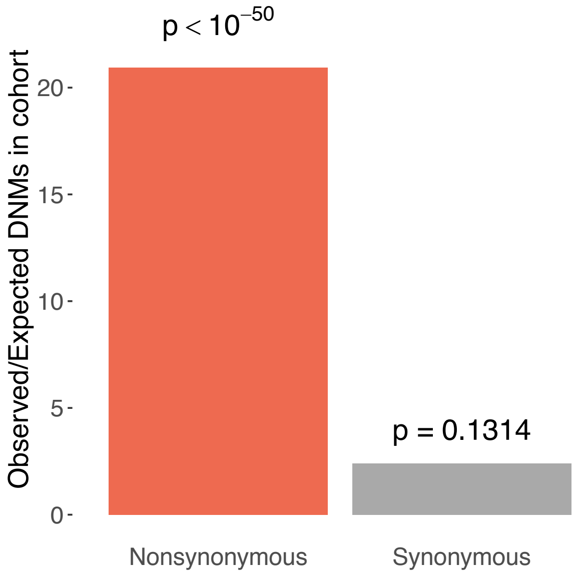


**Supplementary Figure 10**. Enrichment of *de novo* mutations (DNMs) in our cohort of developmental disorder patients at sites that are recurrently mutated in 3 or more patients in The Cancer Genome Atlas (TCGA). Nonsynonymous DNMs (red) are significantly enriched ($p < 10^{-50}$, Poisson test), but synonymous DNMs (grey) are not ($p = 0.1314$, Poisson test).

### DNM burden in non-significant genes

We calculated the remaining DNM burden in the genes that were not significantly associated with developmental disorders (DD) in our analysis and were not a consensus DD gene. There were 2,504 genes that were not associated with DD and had a pLI ≥ 0.9 (high pLI) and 15,413 genes with a pLI < 0.9 (low pLI)[36]. The burden was calculated by calculating the observed/expected number of DNMs across the four groups of genes categorised by both missense/PTV mutations and low/high pLI. While there was a remaining burden across all groups, we found that the highest remaining enrichment was in PTV DNMs in high pLI genes. We then repeated the analysis removing nominally significant genes (unadjusted p-value > 0.01). We saw that the PTV enrichment dropped substantially more than the missense enrichment in both low and high pLI genes (**Sup Fig 11)**. This suggests that our test is still not as well positioned to identify genes with mutations acting via gain-of-function mechanisms that tend to be driven by missense mutations.

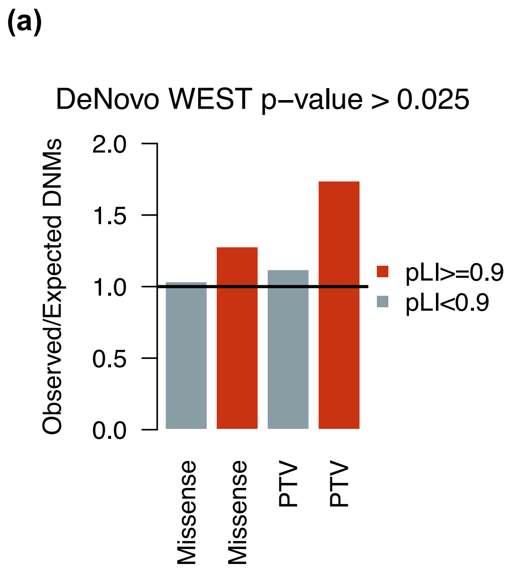


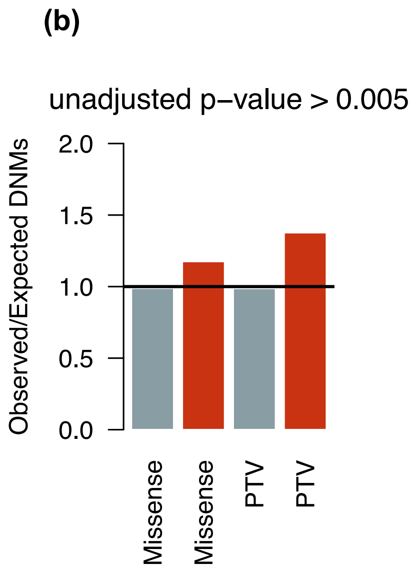


**Supplementary Figure 11: DNM enrichment in non-significant genes.** (a) The enrichment of missense and PTVs separated by high pLI (red) and low pLI (blue). Unassociated genes is defined as those not on any diagnostic list (not consensus or discordant) and not significant in our analysis. (b) The same as (a) except here the genes have been subsetted further to those that are not nominally significant in our analysis (unadjusted p-value <0.01)

### Impact of pre/perinatal death on power

A structural malformation (detected via ultrasound) during pregnancy is associated with pre/perinatal death, which here encompasses spontaneous fetal loss, stillbirth, early neonatal death, and termination of pregnancy for fetal anomaly. To understand the impact that pre/perinatal death may have had on our power to detect DD-associated genes we asked a clinician to assign each consensus known DD gene (n=331) a low, medium or high likelihood of a patient with a pathogenic mutation in the gene presenting with a structural malformation on ultrasound. We subsetted these to those known to be haploinsufficient according to DDG2P (n= 189).We then calculated the ratio of the number of observed *de novo* PTVs in each group to the total number of expected *de novo* PTVs across each group. We found that genes with a medium/high likelihood were significantly less PTV enriched in our cohort compared to genes with a low likelihood (p =0.0001, Poisson test). The difference between medium and low was significantly different (p = 0.004, Poisson test) but the difference between low and high was not (p = 0.05, Poisson test).

To verify that the classification was valid, we compared the proportion of individuals in the DDD study with nonsynonymous mutations in the three gene groups that were reported to have an abnormal scan during pregnancy. We found that the proportion of patients with an abnormal ultrasound for those with a nonsynonymous DNM in a gene with a high or medium likelihood of abnormal ultrasound was significantly higher than for those with a DNM in a gene with a low likelihood classification (12.8% (low) vs 30.0% (medium/high), $p = 9.75 \times 10^{-13}$ ). We also looked at DNMs called in 640 trios from the Prenatal Assessment of Genomes and Exomes (PAGE)[58] study and found that there was a significantly higher enrichment of non-synonymous DNMs in genes with a medium/high likelihood of presenting with an ultrasound abnormality compared to the genes with a low likelihood (2.4 (low) vs 5.0 (medium/high) enrichment, p = 0.049, Poisson test).

For the DDD data, we also had information on whether there had been an abnormal ultrasound during pregnancy for the patients. We regressed the proportion of individuals with a nonsynonymous mutation in a consensus known HI gene who had an abnormal ultrasound against the observed PTV enrichment in that gene. We performed a generalized linear model with a quasipoisson family on the count of PTVs per genes with the expected count and the proportion of abnormal ultrasounds as independent variables. We did not find a significant regression coefficient for proportion of abnormal ultrasounds (p = 0.09).

To assess whether mutations in novel genes are more likely to be associated with pre/perinatal death than consensus known genes we compared the nonsynonymous *de novo* enrichment of these two groups in PAGE. We did not find a significant difference between the enrichment in novel genes (2.6) compared to consensus known genes (3.3) ($p = 0.3$, Poisson test). We are not well powered here, and would need to include more samples to detect a significant difference. We also did not find that individuals in DDD with mutations in novel genes were significantly more likely to have an abnormal ultrasound compared to consensus genes ($p = 0.6$, proportion test 0.6).

### Penetrance

To begin to explore potential variable penetrance, we focused on the 172 genes with significant PTV enrichment ($p < 1 \times 10^{-6}$) and extracted their observed PTV and synonymous counts in gnomAD[36] from the released gene-level constraint file (gnomad.v2.1.1.lof_metrics.by_gene.txt.bgz). Since we planned to compare the PTV to synonymous ratio in these genes, we wanted to remove any genes that were outliers in gnomAD or would have higher-than-expected PTV counts due to biological reasons. We therefore removed the 13 genes that had a constraint flag in the gnomAD file (10 with "syn_outlier", 2 with "mis_too_many|syn_outlier", and 1 with "lof_too_many") as well as three genes associated with clonal hematopoiesis of indeterminate potential (*DNMT3A*, *ASXL1*, and *PPM1D*)[59,60] that were in the 172 PTV enriched genes.

The remaining 156 genes were divided into five bins based on their log10(PTV enrichment). Specifically, the bins were equally spaced in log10(PTV enrichment) space, and therefore vary in the number of genes in each bin (range 9 to 62). For each bin, we determined the PTV to synonymous ratio in gnomAD (**Fig 3c**). Then, we regressed the PTV to synonymous ratio on the PTV enrichment bin and weighted it by the number of genes in each bin. The slope of this regression was nominally significant (p = 0.03069).

#### Power analysis

Calculating power tailored specifically according to our *de novo* enrichment test was challenging as we would be required to make assumptions about the distribution of mutation consequences according to undiscovered DD-associated genes. Even with these assumptions, the calculation would be computationally intensive given the simulation based framework. We decided to calculate power in the context of the enrichment of PTV mutations. Therefore, this power measure is specifically the power to detect haploinsufficient genes. Power was defined as the power to detect the median PTV enrichment in known monoallelic DD-associated genes. The set of known monoallelic DD-associated genes was defined as 146 genes in the consensus gene list that had at least one observed *de novo* PTV in our dataset. We calculated the PTV enrichment by dividing the observed number of *de novo* PTVs in each gene by the expected number of *de novo* PTVs as defined by the gene-specific mutation rate. The median of the PTV enrichment distribution was 37.4. Power was then calculated as the probability of observing a significant p-value under the Poisson test if we assume the median PTV enrichment.

#### Downsampling

To assess if our discovery of DD-associated genes via DNM enrichment has begun to plateau, we downsampled the cohort to 5k, 10k, 15k, 20k and 25k and ran the DeNovoWEST framework to identify the number of significant genes. As is depicted in **Fig 4a**, we do not see evidence of a plateau, indicating there is still much to be gained from DNM enrichment analyses.

### Modelling remaining DNM burden

We modelled the remaining DNM burden separately for PTV and missense mutations.

#### Model for PTV DNM burden

We simulated the number of *de novo* PTVs across a range of numbers of remaining haploinsufficient genes and the PTV enrichment we would observe in these genes. Our PTV model made the following assumptions:

- PTV enrichment was the same across all remaining undiscovered HI DD-associated genes.
- All undiscovered HI DD-associated genes have the same level of penetrance
- The likelihood of being an undiscovered HI DD-associated gene was $1 - power$, where power was calculated as described in a previous section
- The probability of a currently unassociated gene being selected as a HI DD gene was higher if the gene was intolerant of loss-of-function variation. Specifically, the likelihood was multiplied by the observed relative likelihood of being a DD-associated gene (significant in our analysis or a consensus known gene) for genes with pLI ≥ 0.9. This was defined as:

$$\frac{P(DD\ associated\ gene\ |\ pLI \geq 0.9\ \&\ power > 0.8)}{P(DD\ associated\ gene | pLI < 0.9\ \&\ power > 0.8)}$$

- The PTV enrichment for known DD-associated genes (significant in our analysis or consensus known) was taken as the observed enrichment in our cohort
- We ignore missense variants that may act as loss-of-function

To assess the likelihood of the model we calculated the following across the distribution of all 200 simulations per scenario:

$$Likelihood\ = P(6{,}861\ PTV\ DNMs) \times P(147\ significantly\ PTV enriched\ genes) \\ \times P(2{,}929\ genes\ with\ PTV\ enrichment\ > 2)$$

This captures three essential parts of the distribution of observed *de novo* PTVs: the total number of *de novo* PTVs, the number of genes that are currently significantly enriched for PTVs in our cohort and the number of genes that are not significant but have an elevated PTV enrichment ( > 2).

#### Model for missense DNM burden

The set up for the model for missense mutations was very similar to the PTV model with a few key differences. We simulated the number of *de novo* missense variants across a range of numbers of genes with a pathogenic DD-associated variant, mean missense enrichments in these genes and distributions of missense enrichment. We included a third dimension to the likelihood space which allowed us to model the distribution of missense enrichment across the genes with a pathogenic DD-associated missense variant.

We modelled the distribution of missense enrichment using the gamma distribution. We used 6 different shape parameters to represent different scenarios (0.1,0.5,1,510,20) (**Sup Fig 12(a)**). Missense mutations acting via gain-of-function mechanisms are likely to act on a small mutational target and have a small missense enrichment. A shape parameter of 0.1 represents the model that most genes with pathogenic missense DNMs are acting via gain-of-function and have smaller missense enrichment values. Missense mutations acting via loss-of-function mechanisms in haploinsufficient genes will have a distribution similar to PTV enrichment values which tend to be larger. A shape parameter of 20 represents the model where most genes with pathogenic missense variants are acting via loss-of-function and have larger PTV enrichments. The other major difference from the PTV model was that the likelihood of being selected as a DD-associated gene in the simulations was not a function of pLI. It was only proportional to $1 - power$.

To assess the likelihood of the model we calculated the following across the distribution of all 200 simulations per scenario:

$$Likelihood = P(27{,}139\ missense\ DNMs) \times P(130\ significantly\ missense\ enriched\ genes) \times P(3{,}764\ genes\ with\ missense\ enrichment\ > 2)$$

We found that the most likely scenario was that the shape for the missense enrichment distribution was 20. According to this shape there are ~1000 genes with pathogenic DD missense mutations with ~2 fold enrichment of missense mutations. This result complements the implications of the PTV model and suggests that the majority of the remaining pathogenic missense mutations are acting via haploinsufficiency **Sup Fig 12(b)**.

(a)

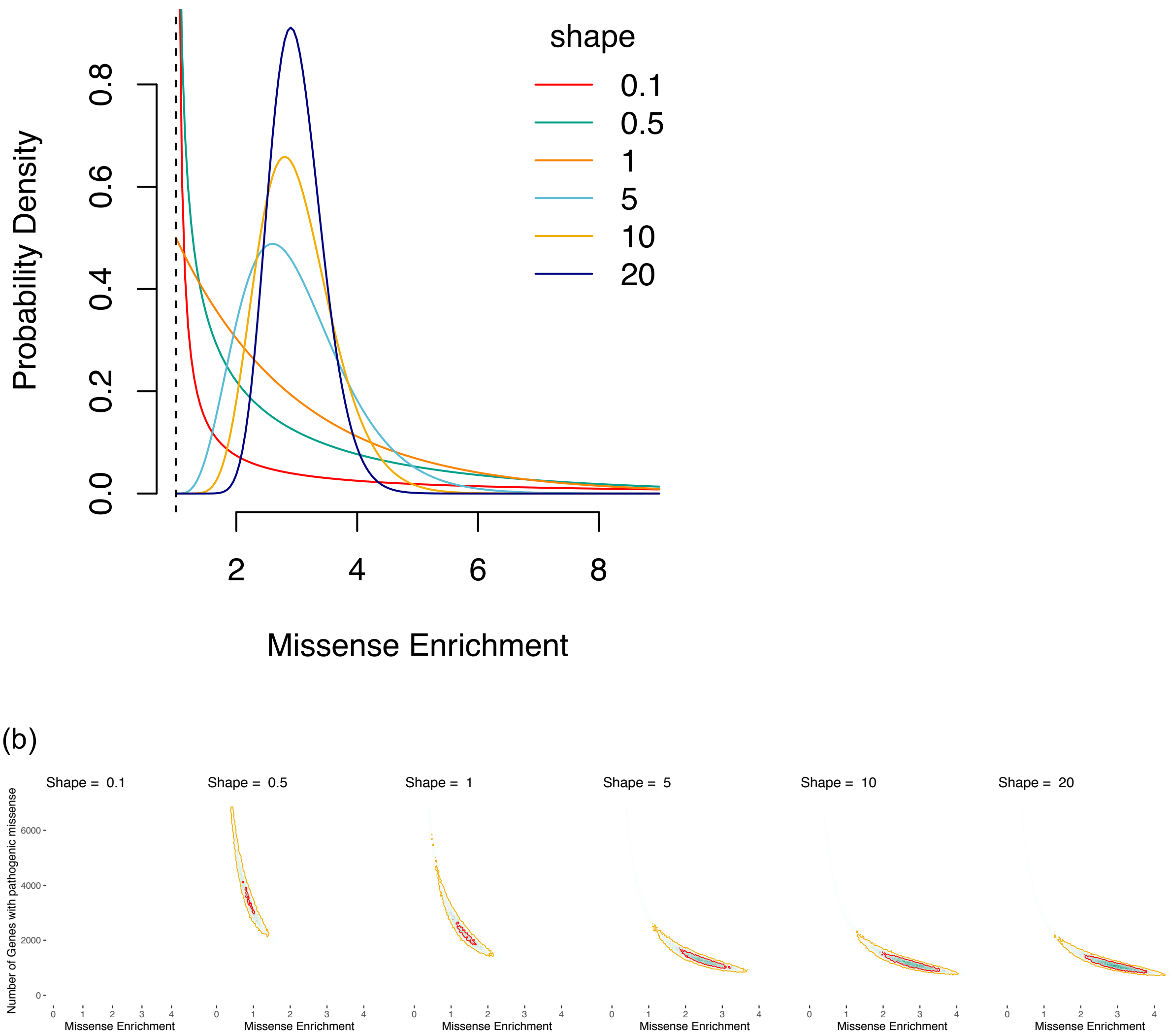


**Supplementary Figure 12**: (a) Depiction of gamma distribution for 4 shape values used in simulations. Here the mean of each distribution is set at 2. (b) Likelihood of scenario under variety of shapes for gamma distribution and under varying values of missense enrichment and number of genes with pathogenic missense variants. The 90% (yellow line) and 50% (red line) credible intervals are calculated across all shapes considered.

### Expression in fetal brain

We defined expression in the fetal brain for a gene as having a median RPKM > 0 in the BrainSpan dataset[31]. We observed that the genes that were not significant in our analysis were significantly less likely to be expressed in the fetal brain ($p = 1.38 \times 10^{-33}$, proportion test). We then subsetted genes into those with a high pLI (pLI ≥ 0.9). We found that there was no significant difference between the proportion of the unassociated high pLI genes expressed in the fetal brain compared to the significant high pLI genes (p = 0.16 , proportion test; **Sup Fig 13**). This suggests that a number of these high pLI genes might be associated with DD as sample size increases.

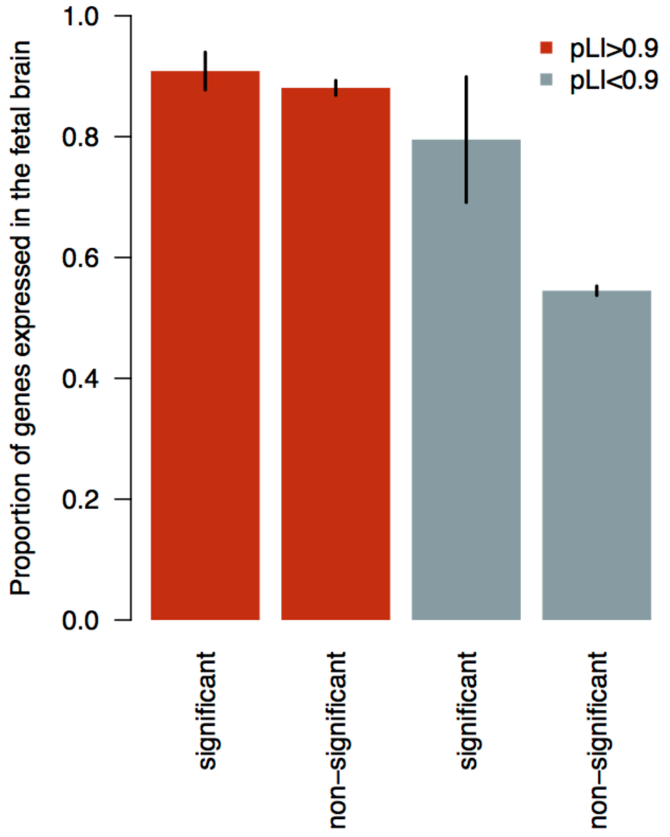


**Supplementary Figure 13: Comparison of proportion of genes expressed in fetal brain.** The proportion of genes between significant and non-significant genes in our analysis split by low (<0.9) and high (>0.9) pLI. Significant genes also includes all genes on the consensus diagnostic list.
